# Bioprinting of aptamer-based programmable bioinks to modulate multiscale microvascular morphogenesis in 4D

**DOI:** 10.1101/2024.06.15.599146

**Authors:** Deepti Rana, Vincent R. Rangel, Prasanna Padmanaban, Vasileios D. Trikalitis, Ajoy Kandar, Hae-Won Kim, Jeroen Rouwkema

## Abstract

Dynamic growth factor presentation influences how individual endothelial cells assemble into complex vascular networks. Here, we developed programmable bioinks that facilitate dynamic VEGF presentation to guide vascular morphogenesis within 3D-bioprinted constructs. We leveraged aptamer’s high affinity for rapid VEGF sequestration in spatially confined regions and utilized aptamer-complementary sequence (CS) hybridization to tune VEGF release kinetics temporally, days after bioprinting. We show that spatial resolution of programmable bioink, combined with CS-triggered VEGF release, significantly influences alignment, organization, and morphogenesis of microvascular networks in bioprinted constructs. The presence of aptamer-tethered VEGF and the generation of instantaneous VEGF gradients upon CS-triggering restricted hierarchical network formation to the printed aptamer regions at all spatial resolutions. Network properties improved as the spatial resolution decreased, with low-resolution designs yielding the highest network properties. Specifically, CS-treated low-resolution designs exhibited significant vascular network remodeling, with increase in vessel density(1.35-fold), branching density(1.54-fold), and average vessel length(2.19-fold) compared to non-treated samples. Our results suggests that CS acts as an external trigger capable of inducing time-controlled changes in network organization and alignment on-demand within spatially localized regions of a bioprinted construct. We envision that these programmable bioinks will open new opportunities for bioengineering functional, hierarchically self-organized vascular networks within engineered tissues.

## 1. Introduction

Three-dimensional (3D) bioprinting of vascularized engineered tissues holds significant promise for creating clinically relevant constructs that closely mimic native tissues, advancing tissue engineering.[1] Despite this potential, *in vivo* outcomes often fall short, particularly in integrating engineered vessels with host vasculature (anastomosis). Human blood vessels exhibit a hierarchical organization facilitating efficient bi-directional transport of oxygen, nutrients and metabolic wastes through capillaries while minimizing energy expenditure. The native vascular hierarchy represents a continuum with gradual tapering vessel sizes as they branch, regulated synergistically by dynamically presented biochemical cues.[2] This hierarchy, essential for proper vascular function, is challenging to replicate in engineered tissues *in vitro*.

Upon implantation, 3D-printed vessels (>400 µm diameter) facilitate vessel retention and perfusion but initially undergo regression in vessel length, diameter, and structural complexity.[3–6] Microvessels (∼50-200 µm) exhibit variable success in terms of anastomosis and perfusion.[2] Direct comparisons between 200 µm and 400 µm diameter 3D-printed vessels within fibrin hydrogels revealed that smaller vessels could not rescue perfusion in a hind-limb ischemia model after 21 days post-implantation.[3] This failure of bioprinted constructs *in vivo* can partly be attributed to the lack of control over *in vivo* remodeling processes and the design’s inability to mimic the multiscale, hierarchical organization of native blood vessels. To achieve effective molecular transport throughout the entire thickness of engineered tissues within the diffusion limitations of ∼200 µm[7] and ensure long-term functionality, it is imperative to incorporate multiscale hierarchical vasculature.

Vascular morphogenesis involves a series of processes where cells dynamically interact with the extracellular matrix (ECM), acting as a dynamic biomolecular scaffold. Processes like vasculogenesis (de novo vessel formation), angiogenesis (new vessel formation from existing ones), vessel assembly, maturation and remodelling are regulated by various angiogenic factors in a spatiotemporally modulated manner.[8] The ECM composition dictates the mechanical properties of the cell microenvironment, including stiffness and viscoelasticity, and regulates multiple secreted angiogenic factors for guiding vascular network morphogenesis.[8] ECM components sequester angiogenic growth factors (GF), notably vascular endothelial growth factors (VEGF), and present them to cells in a controlled manner.

At the tissue level, VEGFs diffuse through the interstitial space to bind the ECM and cell-surface receptors, creating a VEGF concentration gradient and attracting endothelial sprouts towards hypoxic regions.[8] These gradients are facilitated by VEGF isoforms, each binding with different affinities to VEGF receptors and ECM components. Knockout mice expressing only non-ECM binding VEGFs produced small numbers of leaky, wide-diameter vessels, whereas ECM-binding isoforms formed large numbers of thin, highly branched vessels.[8,9] This highlights the importance of establishing stable concentration gradients within ECM to form functional vasculature *in vivo*. Various angiogenic GFs (VEGF, PDGF, and angiopoietins) function in a space and time-controlled manner at different stages of vascular morphogenesis.[8] Long-range gradients of multiple angiogenic cues, characterized by their interdependent functionality, significantly contribute to the stability of vascular networks,[10,11] emphasizing the importance of time-controlled presentation of angiogenic factors to yield functional and stable vascular networks within engineered tissues.

Recent progress in bioprinting technology has focused on augmenting resolution and printing speed, greatly broadening our capability to fabricate engineered tissues.[12,13] Most bioinks have been engineered with an emphasis on printability and mimicking the dynamic viscoelastic behavior of living tissues,[14–16] yet they frequently overlook the dynamic availability of angiogenic factors during vascular morphogenesis *in vivo*. While several studies have effectively showcased the formation of patterned, perfusable vasculature within 3D-bioprinted constructs,[3–6,17,18] many methodologies inadequately address the dynamic remodeling of vasculature over time. Cellular migration from their initial position within the scaffold post-biofabrication contributes to vascular network remodelling, potentially diminishing network functionality over time.

Given that vascular morphogenesis (network formation and remodelling) is governed not solely by the temporal presentation of angiogenic GF but also by their spatial gradients, advanced bioinks are needed to more accurately mimic the dynamic presentation of GF akin to native ECM. Many angiogenic GFs are sequestered within ECM via affinity interactions and released as required through proteases/MMPs, establishing spatial morphogen gradients in a time-dependent manner that dictates cell migration. This contrasts with traditional bioinks employed for angiogenic GF delivery, which primarily focus on physical entrapment, immobilization, or covalent conjugation of GF within polymeric matrix via degradable linkers or microcarrier encapsulation.[19,20] Such approaches typically rely on passive GF release via polymer degradation or proteolytic cleavage, exhibiting sustained release profiles.[19] Recent work has explored non-specific triggers like light, heparin, ultrasound, or nanomaterials to achieve triggered GF release on demand from bioprinted constructs.[18,21–24] While these triggers enable on-demand GF delivery, they often result in initial burst releases, polymer degradation, and sustained release profile upon triggering, making GF available to tissues for extended durations. Additionally, these approaches encounter challenges with loading multiple GFs and controlling individual release profiles. These limitations underscore the incapacity of current bioinks to adjust GF release rates according to the spatiotemporally changing requirements of growing engineered vasculature. Altogether, this underscores the imperative to design advanced bioinks capable of providing high printability alongside dynamic control over GF presentation to mimic the complexities of the tissue microenvironment.

We propose a programmable bioink system that can dynamically control VEGF presentation with high specificity to guide vascular morphogenesis within 3D-bioprinted engineered tissues. We utilized VEGF_165_-specific aptamers with gelatin methacryloyl (GelMA) to design programmable bioinks. Aptamers are small, single-stranded oligonucleotides that form unique 3D conformations to bind target biomolecule with high affinity and specificity. To achieve temporal control with triggered VEGF release kinetics from bioprinted constructs, we leveraged aptamer-complementary sequence (CS) hybridization, which can reverse the aptamer-VEGF binding using CS with higher affinity towards aptamers than VEGF. This enables on-demand, triggered VEGF release for 24hr using CS as an external trigger. This approach stems from our previous works with aptamer-functionalized hydrogels, enabling rapid VEGF sequestration and CS-triggered VEGF release with the same efficiency from hydrogels, irrespective of their culture time.[25,26] We previously showed that spatially heterogenous aptamer distribution within patterned hydrogels facilitates time-controlled spatial VEGF gradients that can direct vascular network self-assembly and orientation.[26] We postulate that the unique combination of bioprinting-associated spatial control over bioink deposition and aptamer-CS hybridization-associated temporal control over free VEGF bioavailability offers a great opportunity to mimic the ECM’s dynamic VEGF presentation properties within engineered tissues. Considering the dynamic control over VEGF presentation and programmable nature of these aptamer-based bioinks, we refer to them as “programmable bioinks” throughout this manuscript.

To explore the full potential of aptamer-based approaches, it is important to investigate the spatial resolution of aptamer-tethered VEGF and freely diffusing released VEGF within programmable bioinks in guiding vascular network organization and orientation. Moreover, by exploiting the temporal flexibility over aptamer-tethered and free VEGF bioavailability within programmable bioinks using a highly specific external trigger, we can perform *in situ* manipulation of vascular network remodelling within bioprinted constructs, even days after fabrication. To our knowledge, there are currently no cell-laden programmable bioinks reported in the literature that possess aptamers for dynamic VEGF presentation *in situ* and leverage their unique capabilities for guiding vascular network formation, orientation and organization remodelling within 3D-bioprinted constructs.

## 2. Results and Discussion

### 2.1 *Designing aptamers-based programmable bioink with high printability and stable* aptamer-CS molecular recognition, post 3D-bioprinting

To design printable, programmable bioinks, we modified GelMA with VEGF_165_ specific, 5’-acrydite functionalized aptamers via visible-light photoinitiated (ruthenium/sodium persulfate, Ru/SPS) free-radical polymerization reaction. Upon initiation, covalently-crosslinked GelMA polymeric networks are formed that facilitate covalent incorporation of aptamers, via the unsaturated acrydite group, into the network (Fig. 1A). GelMA, with its cell-binding peptides (RGDs), matrix-remodelling moieties (MMPs), and tunable viscoelastic properties, is a promising bioink.[27] Yet, balancing its printability and biological functionality is challenging. High-concentration GelMA bioinks (10-20%, >80% methacrylation degree) offers excellent printability and shape-fidelity, but struggle with cell viability.[28] On the other hand, low-concentration GelMA bioinks (<5%), either pristine or with fillers, maintain cell viability,[29] but lack mechanical strength, impacting processability. To address this, we employed 5% GelMA with medium methacryloyl substitution (∼60%). However, extruding the GelMA/aptamer pre-polymer solution at low-concentration, remains challenging due to its low viscosity and slow gelation rate. To enhance printability, we resorted to a sequential crosslinking approach (physical gelation followed by covalent crosslinking). This method leverages GelMA’s physical gelation properties and has previously shown improvements in flow behavior and extrudability of low-concentration GelMA bioinks (3-7%, ∼80% methacryloyl substitution).[30] The process involved filling GelMA/aptamer/photoinitiators pre-polymer solution into syringes and cooling at 4°C for 20 min. Subsequently, the syringes were used directly for bioprinting on a temperature-controlled bed set at 4°C, followed by 2 min of post-print photocrosslinking. The sequential crosslinking enhances physical crosslinking among GelMA chains, eliminating the need for rapid chemical crosslinking upon extrusion, making it suitable for cell encapsulation and maintaining shape fidelity. Additionally, the low-concentration bioinks would render higher porosity and lower stiffness to the bioprinted construct.[30]

**Figure 1.**
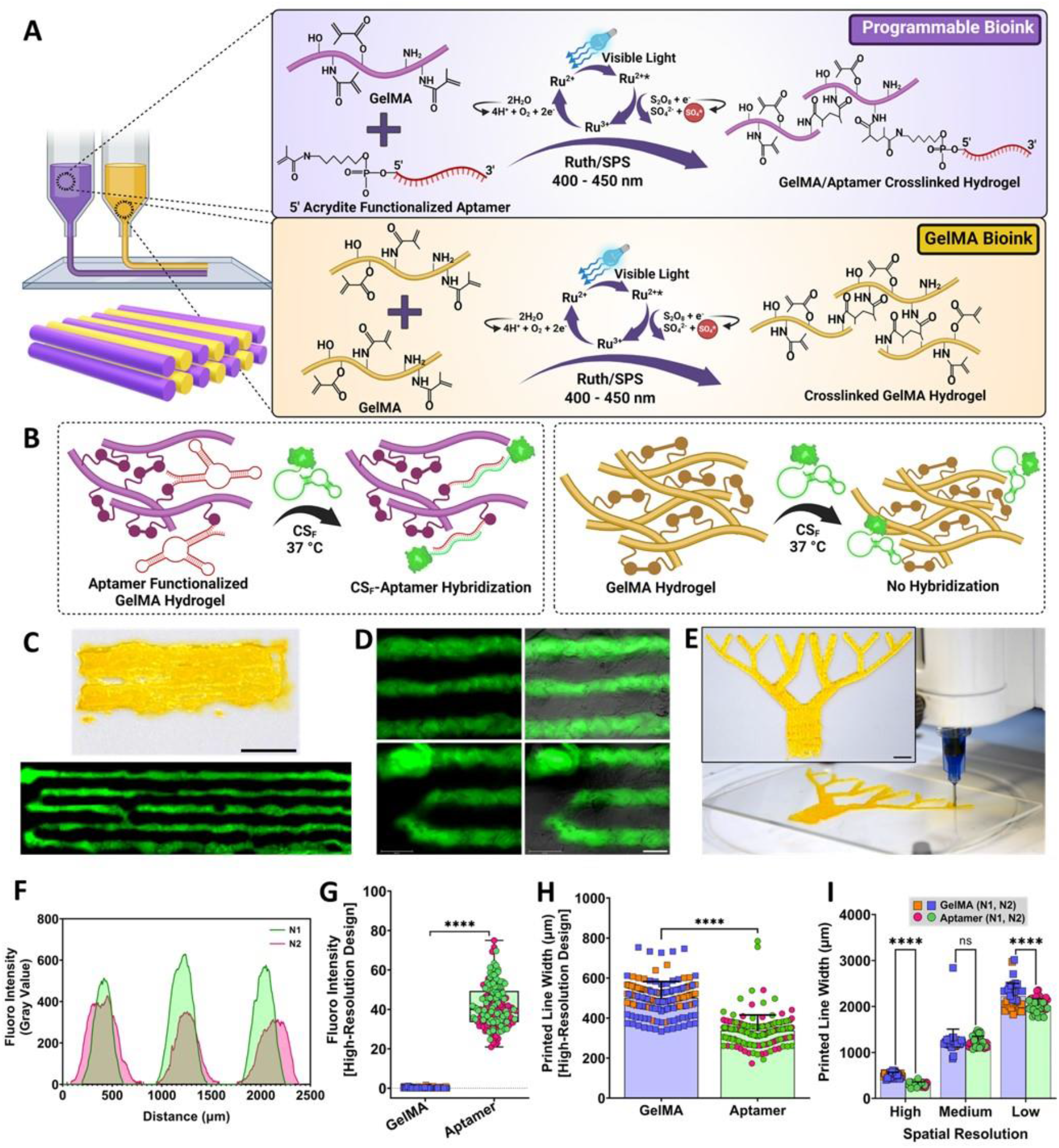
Aptamer-functionalized Programmable Bioink. (A) Schematic representation of aptamer-functionalized programmable bioinks that leverages affinity interaction for controlling GF’s triggered release, alongside pristine GelMA bioink. Upon visible-light exposure, the bioinks with Ru/SPS photocrosslinks the bioprinted constructs. (B, C & D) To validate aptamer retention and aptamer-CS hybridization post-bioprinting, the constructs were incubated with CS_F_ at 37°C for 24hr. The aptamers present in programmable bioink hybridizes with CS_F_, imparting stable fluorescence, whereas, in GelMA regions, the CS_F_ diffuses out of the matrix. (C) Top: Macroscopic photograph of 3D-bioprinted line design. The scale bar is 5mm. Bottom: Fluorescent microscope stitched image of CS_F_-incubated bioprinted design (high-resolution) using programmable and GelMA bioinks, (D) at 10x magnification. The scale bar is 500 µm. (E) Macroscopic photograph of 3D-bioprinted vascular tree design using programmable bioink. The inset scale bar is 5mm. The CS_F-_incubated high-resolution designs after 24hrs were analyzed using two individual samples, represented as N1 & N2, or different colors within same group, for (F, G) fluorescence intensity and (H, I) printed line width. (F) Fluorescence intensity as a function of distance within bioprinted samples, confirms the design consistency. (G) Fluorescence intensity and (H) printed line width among both, aptamer and GelMA regions of the printed construct. The representative microscopic images used for the analysis are shown in (D). (I) 3D-bioprinted line/region width comparison among varying spatial resolution designs (high, medium & low).

Firstly, we printed simple line designs using both programmable and pristine GelMA (i.e., without aptamers) bioinks as continuous filaments, with each bioink printed adjacent to the other followed by photocrosslinking (high-resolution design, 20×6×0.8 mm, Video S1). For the ease of readers, the printed areas with programmable and GelMA bioinks will hereafter be referred as “aptamer regions (A)” and “GelMA regions (G)”, respectively. To assess programmable bioink’s functionality, we first set out to validate the aptamer-CS molecular recognition ability and their retention within bioprinted constructs. To do so, we incubated 5’-Alexa Fluor 488 fluorophore labelled CS (CS_F_) with the printed constructs for 24hr at physiological temperature (37°C), followed by thorough washing with PBS to remove excess/unbound molecules. As anticipated, the printed aptamer regions exhibited significantly higher fluorescence across the construct compared to the GelMA region, attributable to the aptamer-CS_F_ affinity-based hybridization (Fig. 1B,C,D&G). Fluorescence measurements across the cross-section of the bioprinted construct (n=2) revealed overlapping fluorescent regions with nearly similar line widths (Fig. 1F), confirming the programmable bioink’s ability for stable CS_F_ sequestration from the culture medium into the polymeric network, post-printing aptamer-CS_F_ functional molecular recognition, and the reproducibility of printing with programmable bioinks. Additionally, we observed negligible impact of the low-temperature exposure during the physical crosslinking step on the aptamer’s functionality at 37°C, underscoring the critical importance of the aptamer’s conformational stability for its molecular recognition behavior and overall performance. During bioprinting, the bioink being extruded via the nozzle experiences high shear stresses, which can possibly induce conformational changes in aptamers, hampering its functionality.[31,32] However, the observed high fluorescence within printed programmable bioink provides evidence of the chemically conjugated aptamer’s robust stability within the printed polymeric network.

Next, we evaluated the programmable bioink’s print fidelity and reproducibility. Quantification of printed line width in the bioprinted construct revealed that programmable bioinks exhibited significantly higher printing resolution (measured as the printed filament width) (337.8±77.77 µm; coefficient of variance, CV=23.02%) compared to GelMA bioinks (501±81.34 µm, CV=16.24%; p<0.0001), while maintaining constant polymer concentration (5%), printing speed (10mm/s), and nozzle diameter (435 µm) (Fig. 1H). To assess the bioink’s robustness, we progressively increased the design complexity by printing three lines of programmable bioinks adjacent to GelMA bioink (medium-resolution design), followed by a similar pattern for low-resolution designs, where five lines of each bioink were printed consecutively. Additionally, the programmable bioink was utilized to 3D-bioprint a complex vascular tree design with various angles (Fig. 1E). Given their large size, all designs were quantified for their properties using two individual 3D-printed constructs as experimental replicates (n=2). Comparison of the printed line/region widths for the respective bioinks among all designs revealed a consistent trend of significantly higher printing resolution with programmable bioinks, particularly evident in high and low-resolution designs (A-2014±155 µm, CV=7.69%; G-2223.24±278.9 µm, CV=12.55%; p<0.0001), with the exception of the medium-resolution design (A-1233±118.9 µm, CV=9.64%; G-1231±274.7 µm, CV=22.33%) (Fig. 1H&I).

Recognizing that aptamer’s covalent incorporation into the GelMA polymeric network enhances storage modulus of the bulk hydrogel, independent of aptamer amount,[25] we next investigated programmable bioink’s rheological properties. To accomplish this, we photocrosslinked all bioink formulations for 2 min followed by 24hr immersion in PBS to achieve swelling equilibrium. To identify distinct boundaries between the two bioinks within a printed cell-laden construct, we mixed blue fluorescent microbeads (diameter = 2 µm) into the GelMA bioink. Therefore, we performed time-sweep measurements for pristine GelMA, GelMA with blue microbeads and programmable bioinks, as well as amplitude & frequency sweep analysis with their respective crosslinked hydrogels. The frequency sweeps of all samples revealed a significantly higher storage modulus plateau, indicative of ink stiffness, in aptamer-functionalized programmable hydrogels (8897±13.03 Pa, CV=0.15%) compared to the other two samples (Fig. 2B&C). Interestingly, both GelMA and GelMA with microbead hydrogel samples showed storage moduli within a similar range (6133±9.85 Pa, CV=0.16% & 6012±5.77 Pa, CV=0.09% p<0.0001, respectively), highlighting the negligible effect of microbeads mixing on GelMA bioink’s viscoelastic properties at a low particle volume fraction (<0.1). All samples exhibited shear-thinning properties when analyzed for complex viscosity, as evidenced by linearly decreasing complex viscosity with increasing angular frequency (Fig. 2D). However, programmable bioinks exhibited slightly higher complex viscosity compared to other GelMA bioinks. Time sweeps during the photocrosslinking process revealed variations in storage modulus (G’) and loss modulus (G”) among all bioinks as a function of crosslinking time (Fig. 2E). At the onset of crosslinking, while all bioinks showed similar gelation patterns, distinct variations were observed among their modulus profiles. Consistent with frequency sweep results, the programmable bioink exhibited the highest storage modulus (8878±94.95 Pa, CV=1.07%) among all samples, while both GelMA bioinks (GelMA-6234±81.03 Pa, CV=1.30%; GelMA+microbead-6103±54.79 Pa, CV=0.89%, p=0.0010) displayed a similar storage modulus. The enhanced modulus and complex viscosity (better shear-thinning behavior) of programmable bioinks can be directly attributed to the observed higher print fidelity, wherein aptamer’s covalent incorporation renders these bioinks more extrudable. Moreover, aptamers (particularly when present in high amounts) are hydrophilic in nature, which can increase the hydrogel’s swelling properties, resulting in a higher Young’s modulus and larger pore sizes.[25] Based on these findings, we speculate that aptamer covalent incorporation could possibly contribute to the physical crosslinking of GelMA polymeric chains, leading to higher physical crosslink formation within the GelMA network. The subsequent photocrosslinking step stabilizes these crosslinks and generates new chemical crosslinks, further enhancing print fidelity and overall modulus of programmable bioinks.

**Figure 2.**
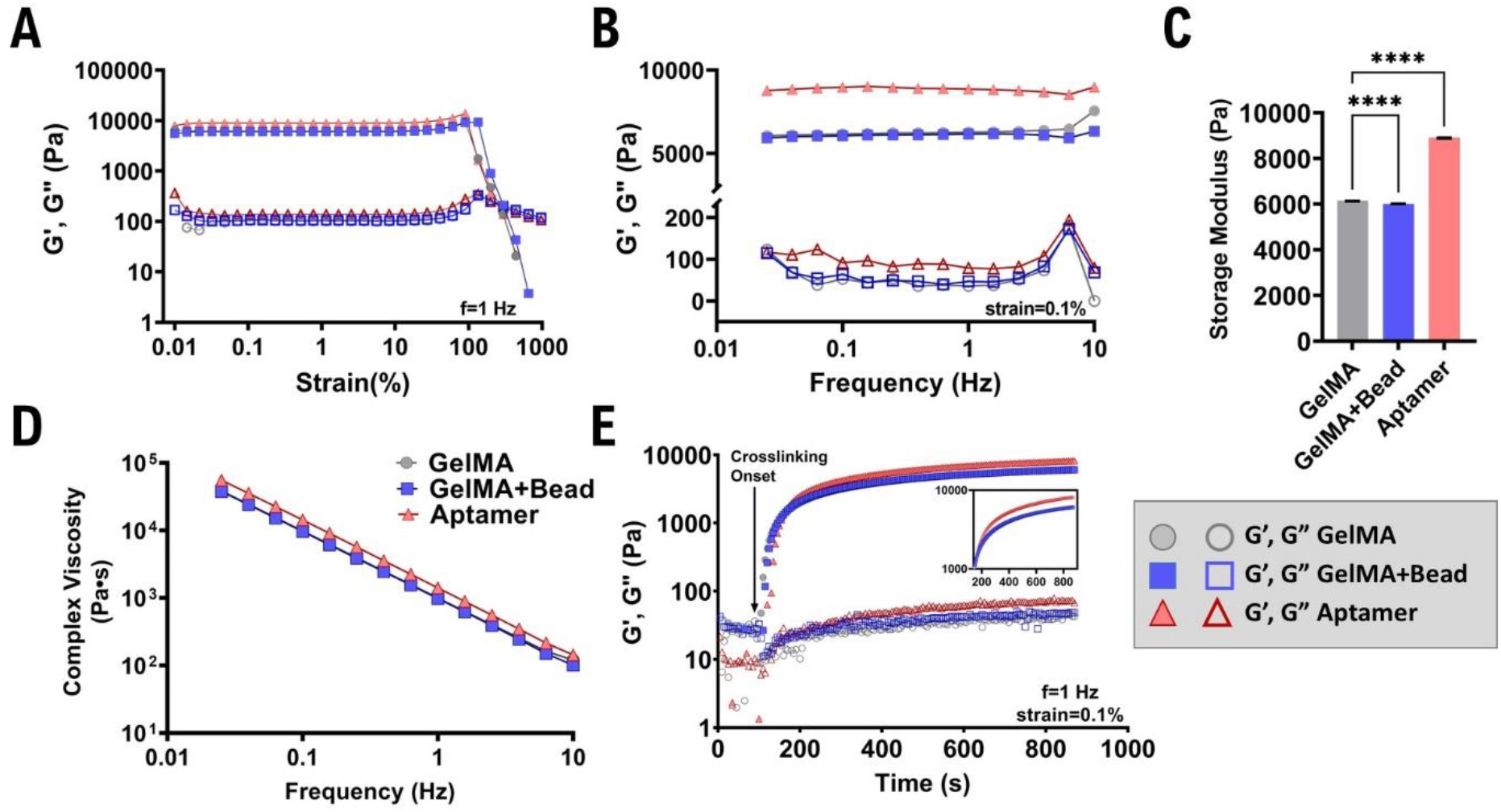
Rheological analysis of the programmable bioink. Rheological analysis of pristine GelMA, GelMA with blue microbeads and aptamer-based programmable bioinks. The storage modulus (filled symbols) and loss modulus (open symbols) of the respective bioinks are shown as (A) strain amplitude sweep at fixed frequency of 1 Hz, and (B) frequency sweep within the linear viscoelastic range at fixed strain % of 0.1. (C) Storage modulus of all bioinks at fixed frequency of 1Hz. (D) Complex viscosity vs. angular frequency plots highlighting the shear thinning (decreased viscosity with continuously increasing angular frequency, 0-100 s^-1^) behavior of all bioinks. (E) Time sweep measurement of the bioinks performed at room temperature (25°C) shows change in storage and loss modulus upon visible light exposure resulting in polymer crosslinking (indicated with arrow). The inset graph with zoomed data plots displays the difference in storage modulus among GelMA-and programmable bioinks with onset of crosslinking.

### 2.2 *Programmable bioink provides temporal control over VEGF bioavailability within* macroscale 3D-bioprinted constructs

Based on the stable molecular recognition of aptamer-CS_F_ confirmed post-bioprinting, we postulate that programmable bioinks can achieve specific VEGF hybridization. We demonstrated previously that bulk aptamer-functionalized hydrogels, incorporating aptamers covalently into the GelMA network (2.5 nmoles), could sequester up to 46% of VEGF from the culture medium within 1hr of incubation, compared to bulk GelMA hydrogels (33%) (VEGF loading amount=10ng).[25] However, aptamer-functionalized hydrogels photopatterned adjacently to pristine GelMA hydrogels displayed pronounced VEGF sequestration from the culture medium. It is likely that photopatterned aptamer and GelMA regions within the same sample induced competitive VEGF sequestration where affinity-based aptamer-VEGF hybridization dominated over gelatin’s inherent electrostatic attraction for VEGF molecules.

Normalized VEGF-antibody fluorescence analysis, measuring sequestered VEGF amount in patterned regions, revealed only 16% VEGF fluorescence within patterned GelMA regions compared to 100% VEGF fluorescence in patterned aptamer regions.[25,26] Considering the prominent effect of the construct’s design on VEGF sequestration ability, this study employed similar adjacent patterns of programmable bioink next to pristine GelMA bioink. To elucidate the effect of spatial resolution on VEGF localization and their temporal release profiles, three bioprinted designs with varying spatial resolution of the printed regions; high, medium and low-resolution designs, were compared.

Programmable bioinks exhibit high specificity and affinity in binding target biomolecules and release them upon triggering by dissociating the aptamer-biomolecule complex, utilizing CS as an external trigger. We have previously demonstrated that bulk aptamer-functionalized hydrogels (2.5 nmoles) leveraging highly specific affinity interactions, could achieve on-demand VEGF release (∼25pg) within 24hr, employing a 1:1 aptamer-CS mole ratio, resulting in a cumulative release of only 30pg over 10d.[25] Maintaining consistent aptamer amount and aptamer-CS mole ratio, it is inferred that similar triggered VEGF release profiles can be attained from printed programmable bioink within macroscale 3D-bioprinted constructs. Given the macroscale dimensions of these constructs, elucidating the spatiotemporal resolution of free VEGF molecules released from programmable bioink is essential. Therefore, we next investigated the influence of design spatial resolution on released free VEGF molecules and their 2D/3D transport behavior within printed macroscale construct over time, employing reaction-diffusion based computational simulations.

The release of free GFs within aptamer-functionalized hydrogels is governed by a combination of diffusion and reaction-binding kinetics between aptamer-GF.[33] The apparent free VEGF diffusivity within aptamer-functionalized hydrogels has been quantified to be (18±2.2)×10^−10^ cm^2^/s.[34] Consistent with literature findings where bulk aptamer-functionalized hydrogels released negligible free VEGF in the absence of CS,[25] it was assumed that VEGF diffusivity within bioprinted designs without CS addition is zero. Numerical simulations mapped the spatiotemporal concentration profiles of the released free VEGF molecules originating from the printed programmable bioink region upon CS addition (0 h) until d6 (144h) (Fig. 3A&B). These simulated timepoints were strategically chosen to align with the experimental timeline of all cell-laden bioprinted designs, where free VEGF release was triggered on d4 and all samples were cultured until d10. The simulation results demonstrate that upon CS-triggering, all bioprinted designs exhibit similar free VEGF diffusion patterns. Initially (0 h), high concentrations of free molecules present in the aptamer regions gradually diffuse towards the centers of the neighboring GelMA regions (Video S2-S7). This diffusion process leads to the formation of instantaneous concentration gradients at the aptamer-GelMA interface, dependent on the spatial resolution of the printed aptamer/GelMA regions (Fig. 3C&D). Higher spatial resolution of aptamer regions within high-resolution designs (region width assumed in model = 400 µm) results in maximal homogenization of these concentration gradients spanning across both regions within 24hr of CS-triggering, compared to other designs. However, in medium (∼1200 µm) and low-resolution designs (∼2000µm), complete homogenization throughout their regions is not achieved; instead, the effect remains localized at the interface up to 144hr.

**Figure 3.**
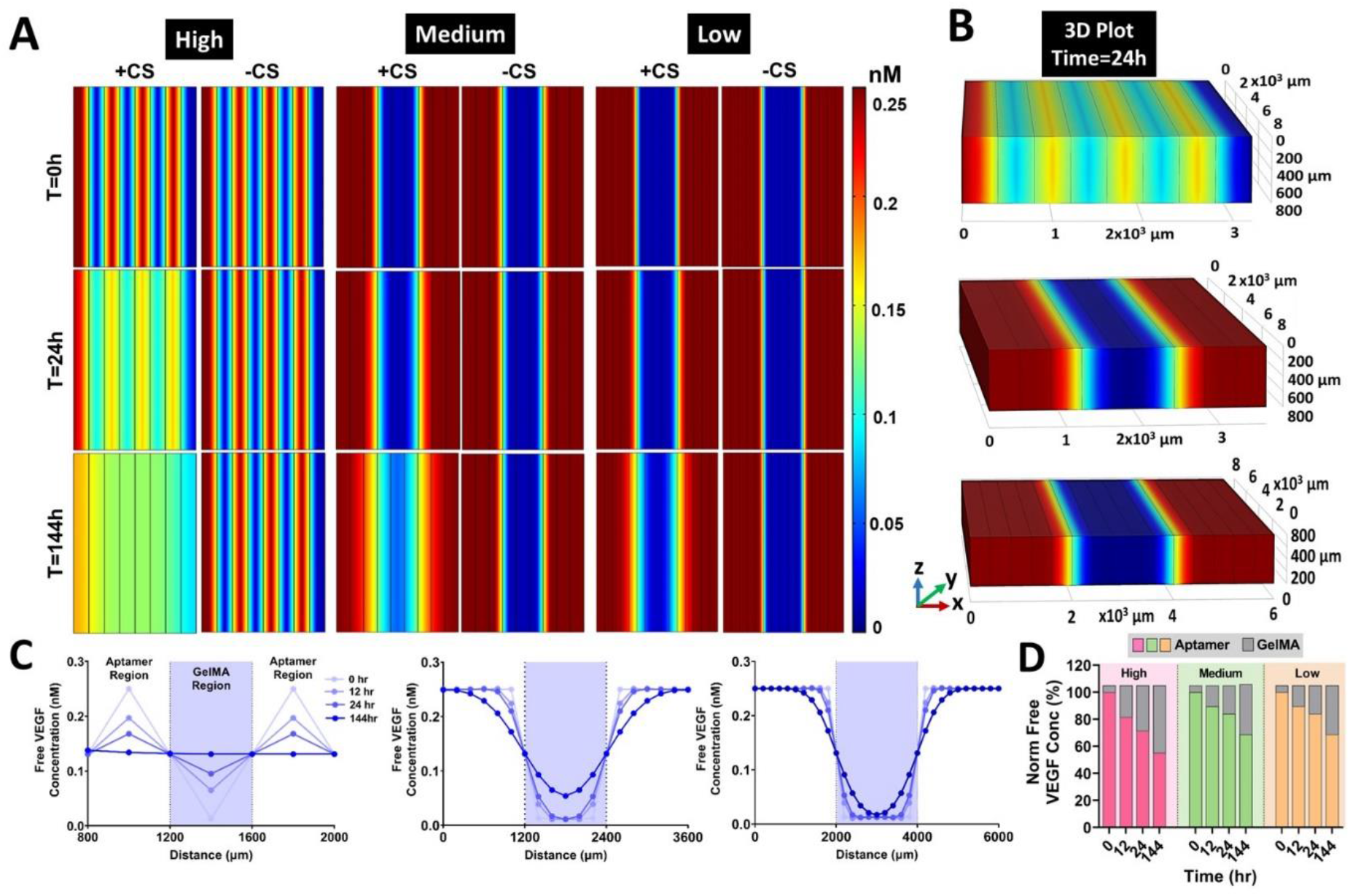
Computational analysis of free VEGF distribution among high, medium and low-spatial resolution 3D-bioprinted designs over time. The (A) 2D and (B) 3D surface plot of free VEGF diffusion from the high (aptamer) to low VEGF concentration regions (GelMA) upon triggered release in the presence (or absence) of CS at different timepoints (t=0, 24 & 144hr). The dimensions of each rectangle among all designs are 400 µm (w) x 8 mm (l). (C) Free VEGF concentration graph as a function of distance across the construct, displaying aptamer and GelMA regions at 0, 12, 24, & 144hr of high, medium & low-resolution designs. (D) Normalized free VEGF concentration% bar graph displaying changes in free VEGF concentrations in both regions over time.

Discrepancies among these designs became more prominent over time, with the free VEGF concentration equilibrating in both aptamer (0.134 nM) and GelMA (0.131 nM) regions in high-resolution designs, while steep concentration gradients persist in medium (A-0.242 nM; G-0.054 nM) and low-resolution designs (A-0.25 nM; G-0.013 nM) 6 days (144hr) after CS-triggering (Fig. 3C). Consistent with the model assumptions, all bioprinted designs showed no free VEGF release in the absence of CS-triggering. The GF diffusivity in the presence of aptamers within a hydrogel is governed by aptamer’s density, binding affinity and matrix pore size.[34,35] This indicates that spatially-controlled, bioprinted constructs using programmable bioink can not only facilitate triggered GF release but also modulate the transport behavior of released free VEGF molecules over time within these constructs.

### 2.3 *Programmable bioink governs cell alignment and shape through spatially defined* VEGF gradients

Our primary goal in designing these programmable bioinks was to model complex processes like vascular morphogenesis in native tissues. Vascular morphogenesis involves dynamic processes such as vasculogenesis, angiogenesis, vessel maturation, and remodeling, all requiring the spatiotemporal regulation of biochemical cues, notably, VEGF. VEGF is a potent angiogenic GF that regulates morphogenesis from embryonic development through adulthood. Inspired by the isoform-specific, heparin-mediated VEGF presentation to cells in native ECM, the aptamers-functionalized hydrogel provides unique opportunity to mimic this spatiotemporally controlled VEGF presentation.

To guide vascular network morphogenesis within 3D-bioprinted constructs, we needed to evaluate the bioactivity of aptamer-tethered and released VEGF molecules within the programmable bioinks. We first assessed the biocompatibility of these 3D-bioprinted constructs where both bioinks were pre-mixed with human umbilical vein-derived endothelial cell (HUVECs) and mesenchymal stromal cells (MSCs) in 1:1 ratio. After printing and photo-crosslinking, the constructs were incubated with medium supplemented with VEGF_165_ (10ng) for 1hr to enable rapid GF sequestration, followed by washing and medium replacement. To ensure that the observed cell behavior was mainly due to the bioavailability of exogenously loaded GF, no VEGF supplements were added into the co-culture medium after the initial VEGF loading step. As a proof-of-concept, we tested bioprinted constructs with three experimental replicates, mixing blue microbeads (diameter=2 µm) with GelMA bioink to easily identify the printed region’s boundary. Fluorescence microscopic images of the bioprinted construct on d1 confirmed homogeneous cell distribution and high cell viability (>90%) in both bioink regions, with most cells exhibiting a round morphology regardless of their region (Fig. S1). These observations confirm the biocompatibility of the bioprinting process, post-print photocrosslinking, and aptamer-mediated VEGF sequestration. Additionally, the presence of cells within the bioinks did not compromise the bioink’s printability when tested for all three designs (Fig. S2).

To determine whether aptamer-mediated VEGF presentation influences cell behavior within bioprinted constructs, we cultured cell-laden constructs using both bioinks and supplemented CS_F_ for 24hr to trigger VEGF release on d4. Fluorescence microscopy revealed notable differences in cell orientation among different regions of the construct, depending on the presence/absence of CS_F_ (Fig. 4). On d5, majority cells in both bioinks displayed elongated morphology and aligned along the printing direction (0° & 180°), consistent with previous reports of MSCs/HUVECs in GelMA-based bioinks.[36,37] In CS_F_-treated samples, cells in both regions showed maximum alignment at 0°-20° (A-18.16%; G-19.60%) and 160°-180° (A-17.97%; G-17.64%). Interestingly, a significant percentage of cells aligned perpendicular to the printing direction (90°) in both regions (80°-100°: A-7.86%; G-9.15%). Non-treated samples displayed similar trends, with the majority of cells aligning at 0°-20° (A-21.56%; G-18.59%) and 160°-180° (A-18.08%; G-17.52%), and a significant number aligning perpendicularly (80°-100°: A-6.10%; G-9.70%). GelMA regions consistently showed higher percentages of perpendicularly aligned cells compared to aptamer regions, regardless of CS_F_-treatment on d5.

**Figure 4.**
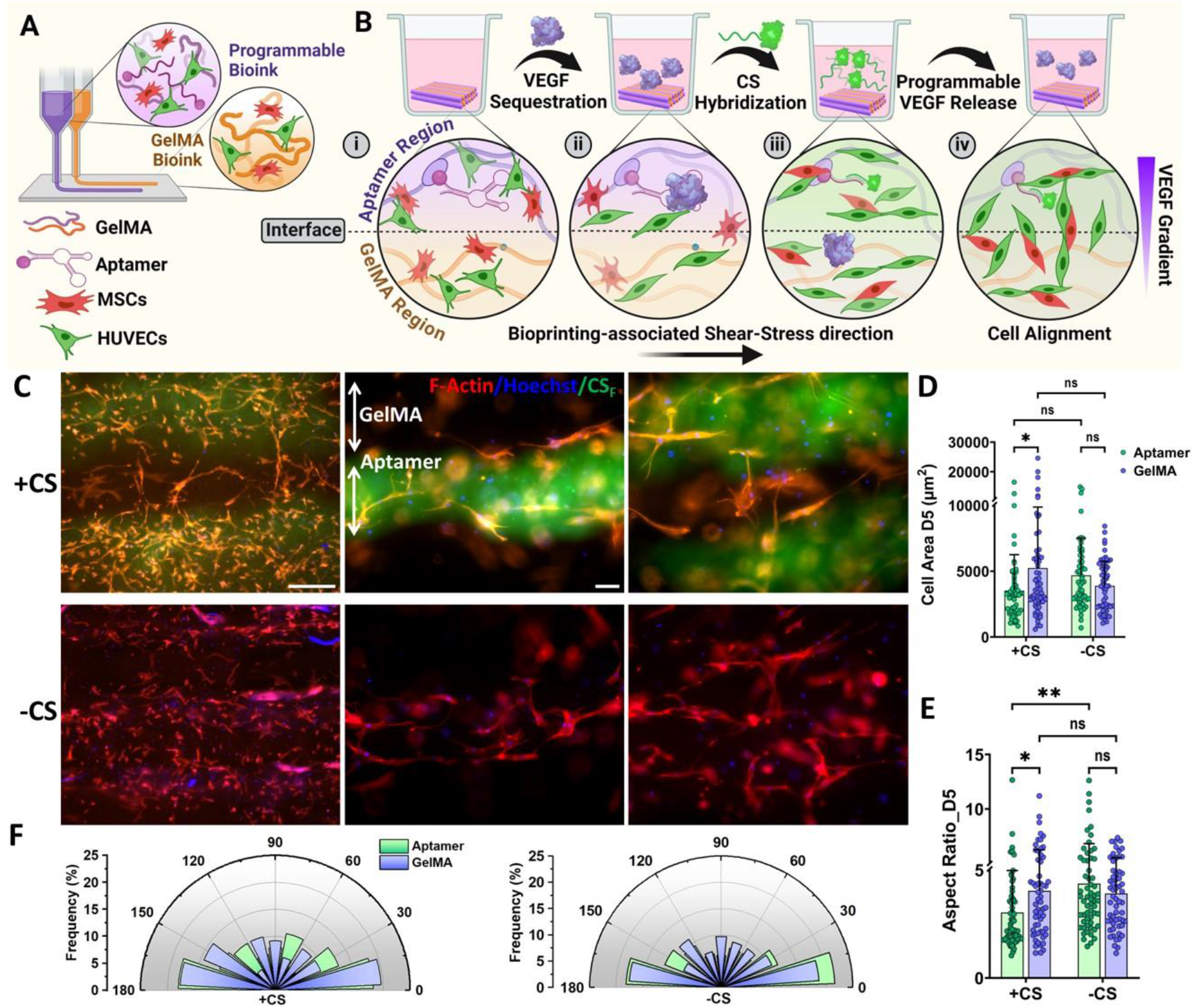
Programmable VEGF bioavailability within 3D-bioprinted constructs selectively guides cell behavior. (A) & (B) The concept of cell-laden programmable bioinks, (i) where aptamer-based programmable bioinks and GelMA bioinks are 3D-bioprinted. (ii) Aptamers facilitate VEGF sequestration from the culture medium to the specific regions in the construct and, (iii) provide programmable VEGF release in the presence of CS_F_, (iv) resulting spatial VEGF gradients within the constructs that guides cell alignment selectively depending on the gradient steepness. (C) F-actin (cell cytoskeleton, red) and Hoechst (nuclei, blue) stained bioprinted samples in the presence/absence of CS_F_ (added on d4) on d5 of culture. The green color corresponds to the presence of CS_F_ hybridized within aptamer regions. The scale bar is 500 µm and 100 µm, respectively. Cellular characteristics such as (D) area, (E) aspect ratio and (F) alignment within aptamer and GelMA regions of the 3D-bioprinted samples on d5 were quantified using ImageJ. The data in (D) & (E) is represented as mean±SD with all data points (n=6, technical replicates). (F) Polar plots representing the cellular alignment and their related frequency (%) on d5 among both regions of bioprinted construct, in the presence/absence of CS (added on d4).

Endothelial cells (EC) are mechanosensitive[38] and experience shear stress during extrusion-bioprinting as the bioink passes through a nozzle.[39] Cells near the nozzle wall experience maximum velocity gradients and shear stress, leading to membrane stretching and deformation.[40] This shear stress induces cell alignment within bioprinted constructs by transforming the actin cytoskeleton, aligning stress fibers in the direction of stress.[41] Consequently, cells elongate and orient parallel to the shear stress, displaying centrally located thick stress fibers.[42] Therefore, the parallel cell alignment observed in bioprinted samples likely results from bioprinting-induced shear stress causing actin cytoskeletalrearrangement.[43]

In this study, cells experienced both biophysical (shear stress) and biochemical (aptamer-tethered VEGF) stimuli simultaneously. Computational simulations showed that aptamer-tethered and released VEGF molecules within bioprinted constructs form spatial VEGF gradients in the presence/absence of CS (Fig. 3). The observed cell alignment along the VEGF gradient direction (perpendicular to the printing direction, 90°) corroborates simulation results. CS-triggered samples exhibited a spatial VEGF gradient from the aptamer region’s midpoint (high concentration) to the GelMA region’s midpoint (low concentration), whereas non-treated samples had a confined VEGF gradient within interface regions. While the precise mechanism of simultaneous shear stress and VEGF sensing remains unclear, it can be inferred that steeper gradients from CS-triggering results in higher cell area (5235.29 µm^2^) and aspect ratio (AR) (4.03) in the GelMA region compared to the aptamer region (area-3518.17 µm^2^; AR-3.03) (Fig. 4D&E). Without CS-triggering, VEGF remains localized in the aptamer region, reversing cell behavior to show the highest area (4684 µm^2^) and AR (4.37) in the aptamer regions rather than GelMA regions (cell area-3889.99 µm^2^; AR-3.91).

ECs exhibit a combined response to fluid-induced shear stress and VEGF-A when both are present at physiological concentrations. ECs align perpendicularly to the direction of flow (shear stress), which differs from the parallel alignment seen without VEGF or with pathologically high VEGF-A levels. The interplay between shear stress and VEGF-A is evident: shear stress alters how sensitive cells are to VEGF-A, and VEGF-A affects how cells respond to shear stress.[44] This behavior corroborates with our observations, suggesting that bioprinted cells sense and respond to the spatial distribution of aptamer-tethered or released VEGF through chemotaxis, overriding bioprinting-associated shear stress. This results in a substantial number of perpendicularly aligned cells in both CS-treated and non-treated samples. The increased cell elongation and area in the GelMA region of CS-treated samples, compared to the aptamer region, likely stem from cells elongating along steeper VEGF concentration gradients through chemotaxis.

### 2.4 *Spatial resolution and CS-triggering synergistically promotes selective cell guidance in* time-controlled manner

To enhanced aptamer-GelMA interface visualization within the bioprinted constructs, blue fluorescent microbeads (2 µm) were added into the GelMA bioink and cell behavior of the constructs were then compared with our previous results from CS_F_-triggered constructs without microbeads, serving as the control on day 5 (Fig. S3). Consistent with our findings from printed samples without microbeads, we observed maximum parallel alignment along the printing directions in samples with microbeads. Regardless of printed regions and CS-treatment, a significant percentage of cells aligned perpendicular to the printing direction (parallel to VEGF gradient) (Fig. S3D&E). However, samples printed without microbeads showed higher cell areas in both aptamer and GelMA regions compared to those with microbeads, in both CS-treated and non-treated samples (Fig. S3A). These differences were statistically insignificant, except in the GelMA region of CS-treated samples (p≤0.0001) (Fig. S3B). Both regions had similar AR in CS-treated and non-treated samples, with statistically insignificant differences, except for the aptamer region of non-treated samples without microbeads (p≤0.05) compared to those with microbeads (Fig. S3C). Microbeads can obstruct cell behavior and affect their response to the microenvironment, depending on their size and density.[45,46] Cells interacting with encapsulated polystyrene microbeads can experience mechano-stimulation due to abrupt mechanical property mismatches. However, in this study, the microbead concentration (1mg/ml) did not significantly affect overall cell behavior. Comparative analysis showed statistically insignificant differences in cell properties such as area, AR, and alignment, regardless of microbead addition. Based on these findings, we included microbeads in subsequent experiments as visual indicators of printed GelMA regions.

Next, to investigate the influence of programmable bioink spatial resolution on selectively guiding cell behavior, we bioprinted cell-laden designs with varying spatial resolution of the printed regions (high, medium & low), loaded with VEGF and cultured till d10, where CS was added on d4. Most cells aligned parallel to the printing direction across all designs, regardless of CS-treatment or culture time (Fig. 5A). The high-resolution design showed the highest cell percentages for various alignment degrees on d5, especially in CS-treated samples in both aptamer and GelMA regions. On d5, non-treated samples exhibited maximum parallel cell alignment followed by perpendicular alignment in the aptamer region. However, in the GelMA region, cells were mostly aligned parallel, with equal distribution in other directions. After 10 days of culture, the majority of cells remained aligned parallel to the printing direction with minimal perpendicular alignment in both regions, regardless of CS-treatment. Non-treated samples had slightly higher cell percentages compared to CS-treated samples on D10 (Fig. 5C). This behavior supports our hypothesis that CS-triggered released VEGF diffuses into the hydrogel and culture medium, remaining available to encapsulated cells in both regions for 24h. On d5, cell alignment in CS-treated samples is influenced by a combination of released VEGF, aptamer-tethered VEGF, and their instantaneous directional concentration gradients within the constructs. This explains the relatively high cell alignment percentages in varied directions among CS-treated samples on d5. This effect was prominent in both regions of high-resolution designs, but decreasing spatial resolution diminished the alignment effect within aptamer regions on d5, becoming more prominent in GelMA regions of CS-treated samples.

**Figure 5.**
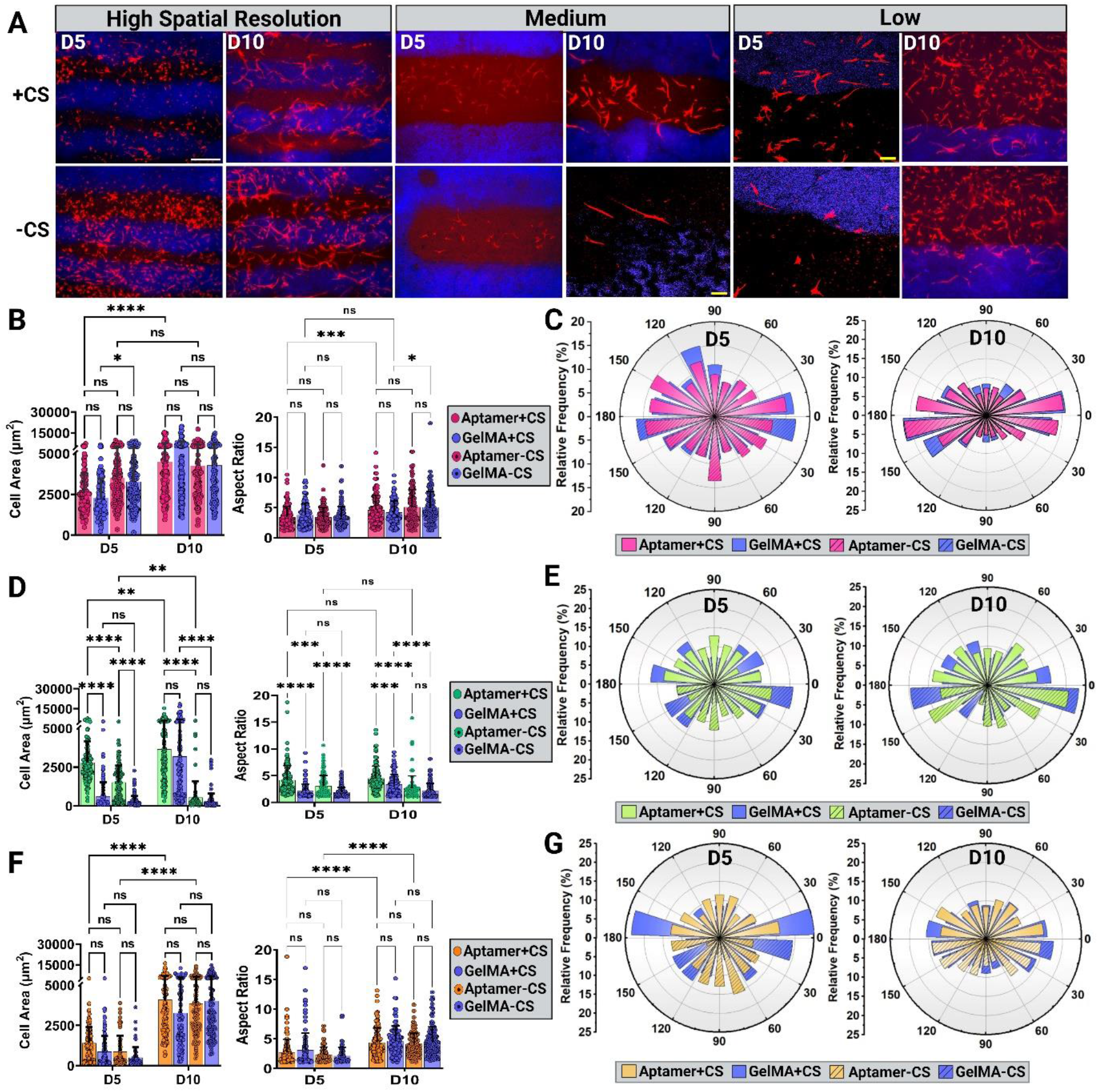
Spatial resolution of programmable bioinks, combined with CS triggering influences printed cell’s characteristics and alignment in time-controlled manner. (A) Fluorescent microscopic images of F-actin (red)-stained bioprinted designs (high, medium & low spatial resolution) in the presence/absence of CS on d5 & d10. Blue color corresponds to the microbeads within GelMA regions. The scale bars are 500µm and 100µm. Cell properties such as area, aspect ratio and alignment on d5 & d10 for (B, C) high, (D, E) medium and (F, G) low-resolution designs, respectively (n=6, technical replicates).

In medium and low-resolution designs on d5, the aptamer regions of CS-treated samples exhibited lower cell percentages aligned in varied directions compared to GelMA regions, where high cell percentages aligned parallel, with the highest observed in low-resolution designs [medium: 0°-20°(A-12.5%; G-12.39%): 160°-180°(A-13.37%; G-16.81%)][low: 0°-20°(A-16.03%; G-25.23%): 160°-180°(A-12.97%; G-23.36%)]. This effect was consistent in non-treated samples on d5 with lower cell percentages (Fig. 5E&G). The high parallel cell alignment within GelMA regions of low-resolution designs can be attributed to the combination of spatial resolution and instantaneous VEGF concentration gradients formed upon CS trigger. As shown in Fig. 3, low-resolution causes steeper VEGF gradients from high concentration aptamer regions to low concentration GelMA regions. EC and MSCs co-cultures are sensitive to VEGF gradients and tend to orient towards high concentrations (via chemotaxis),[47,48] explaining the high cell alignment in GelMA regions of medium and low-resolution designs.

On d10, the low-resolution designs exhibited the least parallel alignment, followed by alignment in varied directions, regardless of region and CS-treatment (Fig. 5G). The medium-resolution designs showed intermediate results, with non-treated samples displaying maximum parallel alignment in GelMA regions followed by perpendicular alignment (Fig. 5E). Overall, the programmable bioink’s spatial resolution, combined with CS-triggered VEGF release, modulates cell alignment over time within bioprinted constructs. Upon CS trigger (temporal control), high-resolution designs yield comparatively random cell alignment in the short term, whereas low-resolution facilitate parallel alignment on d5. However, the CS trigger effect is transient, and over time, cells achieve maximum parallel alignment in high-resolution designs, while cells align randomly in low-resolution designs. This agrees with literature, where high-resolution printed filaments lead to higher cellular alignment.[49,50]

The spatial resolution of the printed construct, combined with CS, influences other cell characteristics like cell area and AR. At high-resolution, cells exhibit high cell areas and ARs on d5, regardless of the region and CS-treatment (Fig. 5B). At medium and low-resolution, high cell areas and ARs are confined to aptamer regions on d5, with notable effects in CS-treated samples compared to non-treated ones (Fig. 5D&F). After 10 days, cells attain higher areas and ARs in all designs, irrespective of CS-treatment, except for non-treated medium-resolution designs (Fig. 5D&F). Although the exact reason for this anomaly is unknown, it may be due to experimental error. These findings suggest that, in the short-term, high-resolution design results in highly elongated cells with large areas, aligned both parallel and perpendicular to the printing direction. Over time, these constructs maintain elongated cell morphology with increased area, with most cells aligning parallel to the printing direction. The effect of CS-treatment is more evident at medium and low-resolution, where CS-treated samples show higher cell elongation and areas in aptamer regions compared to GelMA regions on d5. At low-resolution, CS-triggering promotes parallel cell alignment in GelMA regions on d5, although this alignment effect diminishes over time, with cells maintaining high area and elongated morphology.

### 2.5 *Spatially bioprinted programmable bioink combined with CS-triggering modulates* multiscale EC morphogenesis in 4D

Next, we assessed the influence of spatial resolution on EC morphogenesis within bioprinted constructs. Constructs with high, medium and low-resolution were bioprinted and cultured until d10, with CS added on d4. Confocal microscopy of interface regions revealed that self-assembled CD31+ vascular networks and morphogenesis were confined to the aptamer regions of the constructs, regardless of CS-triggering and culture duration (Fig. 6A). Fluorescence intensity, measured across the interface region, indicated that most fluorescence peaks occurred within the aptamer regions rather than in the GelMA regions. However, in medium and low-resolution designs, a few peaks appeared in the GelMA regions on d10, limited to the immediate vicinity of the interface (∼<300 µm) and not extending across the entire GelMA region (Fig. 6B). Quantification of network morphometric parameters confirmed that the design’s spatial resolution, CS-treatment and time collectively influenced the overall organization of the vascular network.

**Figure 6.**
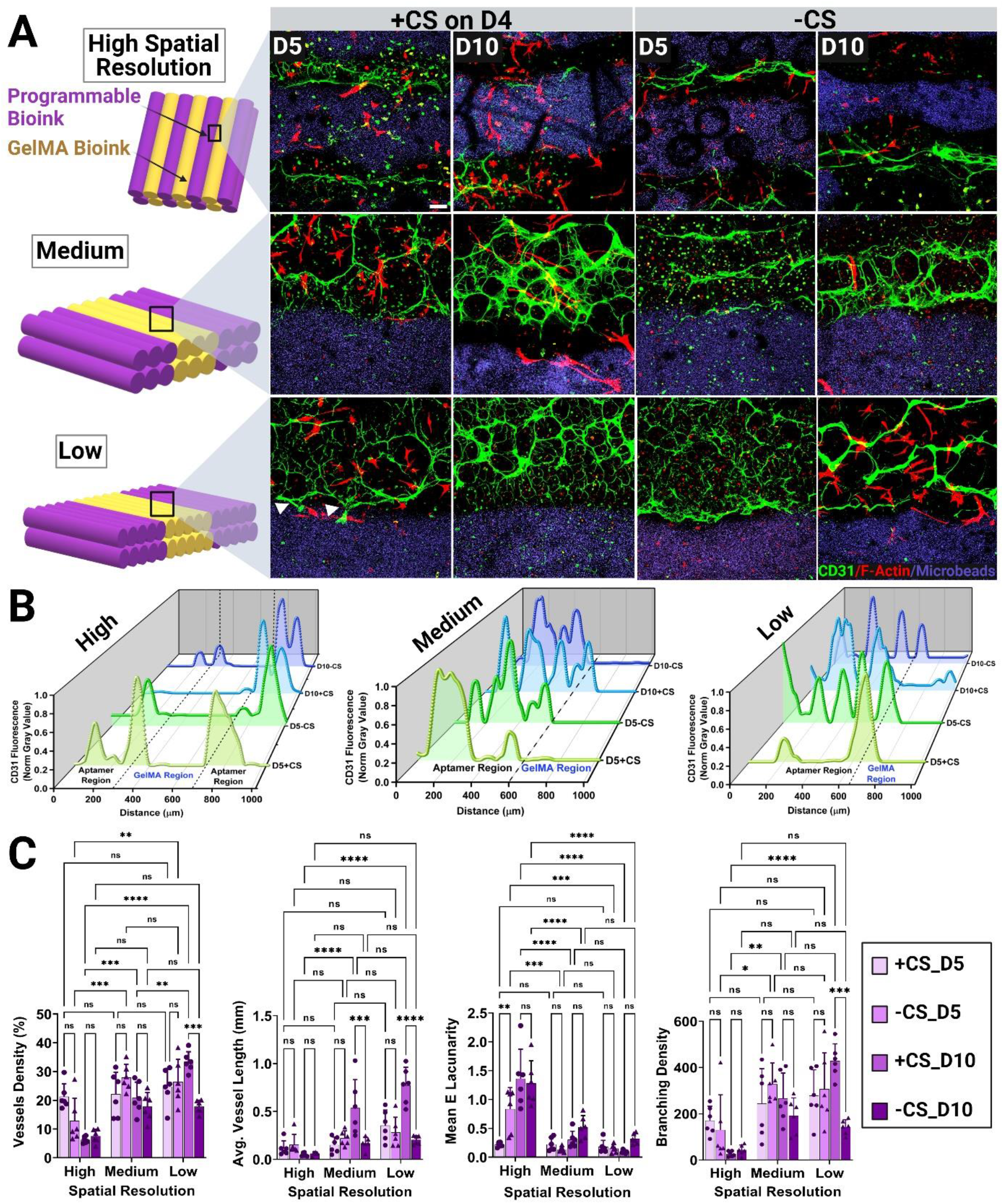
Spatial resolution of programmable bioink combined with CS triggering modulates vascular network self-organization over time. (A) Schematic illustrating high, medium and low-spatial resolution bioprinted designs. Confocal z-stacks of all designs, in the presence/absence of CS (added on d4) on d5 and d10, showing the interface between aptamer and GelMA regions. Blue color corresponds to fluorescent microbeads present within GelMA region. Immunostained samples for cytoskeletal F-actin (red) and EC specific CD31 marker (green). The scale bar is 100 µm. (B) Normalized CD31+ fluorescence intensity over interface region (dotted line) in presence/absence of CS at different timepoints denoted as D5+CS, D5-CS, D10+CS & D10-CS for all designs. (C) CD31+ vascular network properties within aptamer regions of all designs over varying spatial resolutions on d5 & d10. High-resolution corresponds to 340 µm, medium-resolution to 1230 µm and low-resolution to 2014 µm of aptamer region’s width (n=6, technical replicates).

Given that vascular network formation was predominantly confined to aptamer regions, we first compared network properties among aptamer regions of all designs (Fig. 6C&S4). Vessel properties increased as programmable bioink’s spatial resolution decreased, with low-resolution designs yielding the highest vessel properties. CS-treated medium and low-resolution designs showed minor differences in vessel properties on d5, which became significant by d10, indicating the substantial impact of CS-treatment over time. For example, CS-treated samples exhibited significantly higher average vessel lengths on d10 compared to d5 for both medium (d5-0.149 mm; d10-0.536 mm) and low-resolution designs (d5-0.353 mm; d10-0.774 mm). Conversely, average vessel length among non-treated samples remained almost similar at both time-points, regardless of spatial resolution [(medium: d5-0.221 mm; d10-0.167 mm)(low: d5-0.282 mm; d10-0.204 mm)]. Network lacunarity, indicative of nonuniformity, exhibited a linear relation with the spatial resolution, with high-resolution designs showing maximum lacunarity compared to medium and low-resolution designs, regardless of time. Notably, CS-treated samples demonstrated decreased lacunarity compared to non-treated samples on d5, suggesting that CS-triggered VEGF release positively influenced vascular morphogenesis by reducing network nonuniformity. *In vivo*, high lacunarity is characteristics of pathological vasculature, implying that low lacunarity is favorable for healthy vessel organization.[51] This CS effect was more pronounced in high-resolution designs, though other designs followed the same trend [high-resolution lacunarity, d5(+CS: 0.217; –CS: 0.825); d10(+CS: 1.359; –CS: 1.279)]. The lower lacunarity in medium and low-resolution designs resulted from the higher uniformity of vascular networks, with higher vessel and branching densities contributing to this uniformity.

Combining spatial resolution with CS-treatment resulted in three major types of vascular network self-organization: single-vessel-based fibrillar networks (in high-resolution designs), densely packed networks with low intervessel spaces, and hierarchically organized networks with variable intervessel spaces (in medium/low-resolution designs). The intervessel space, referring to the void space between connected vascular networks, showed an inverse relationship with network self-organization, improving over culture time as confirmed by the confocal microscopy images from various regions (interface, aptamer, and GelMA) (Fig. S4). Given the significant role of the printed GelMA region in guiding cell properties, including alignment (Fig. 5), we quantified vessel properties in the printed GelMA region and compared them with aptamer region results (Fig. 7).

**Figure 7.**
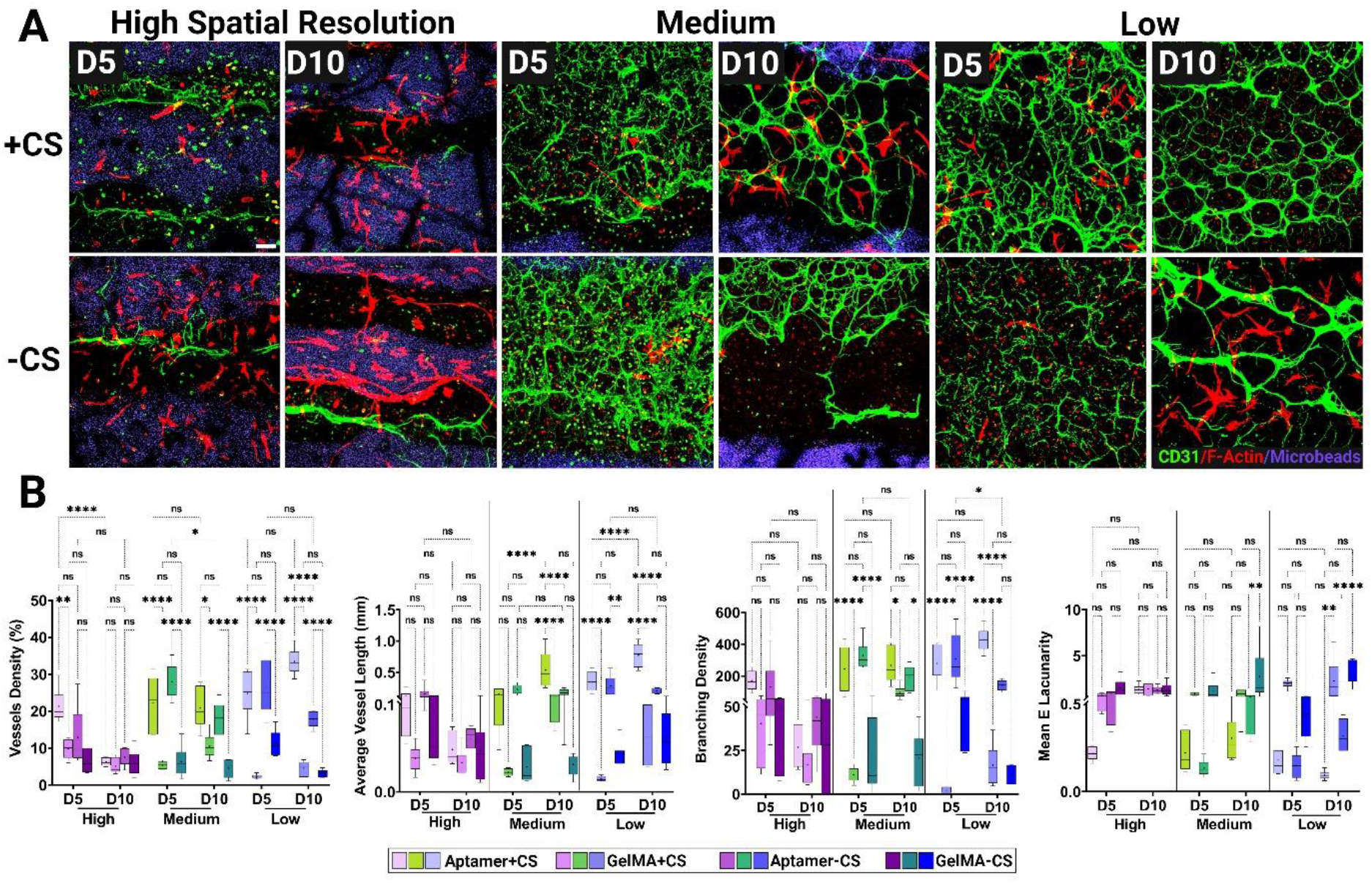
Spatiotemporal localization of aptamer-tethered VEGF in 3D-bioprinted programmable bioink controls endothelial cell morphogenesis confined within aptamer regions. (A) Confocal z-stacks showing cell cytoskeletal F-actin (red) and EC specific CD31 (green) expression in 3D-bioprinted designs (high, medium & low-resolution) in presence/absence of CS (added on d4) on d5 & d10. Blue color corresponds to fluorescent microbeads present in GelMA region. The scale bar is 100 µm. (B) CD31+ vascular network properties in both aptamer and GelMA regions of each design (in the presence/absence of CS) on d5 & d10 (n=6, technical replicates).

CS-treatment immediately impacted vessel organization patterns and properties of CD31+ cells across all experimental designs. Generally, CD31+ cells in printed aptamer regions exhibited greater cell area and AR compared to those in GelMA regions, regardless of spatial resolution and CS-treatment (Fig. S5A,C&E). High-resolution designs showed slight increases in cell properties with CS-triggering, though these differences were statistically insignificant. Notably, CS-treated high-resolution designs featured CD31+ cells with the smallest cell area (1922 µm^2^) and highest AR (3.42) in aptamer regions on d5, which diminished over time (Fig. S5A). The slender, elongated morphology of CD31+ cells in CS-treated, high-resolution designs led to the formation of parallel-aligned, fibrillar-branched vessel networks with higher vessel densities (A-21.31%, G-9.81%) and branching densities (A-170.5, G-40.24) in aptamer regions on d5. In contrast, non-treated samples exhibited longer average vessel lengths [non-treated(A-0.153mm, G-0.071mm); CS-treated(A-0.116mm, G-0.382mm)] with lower vessel densities (A-12.85%, G-6.43%) and branching densities (A-130.7, G-32.97). Over time, CS-treated high-resolution designs showed vessel regression with reduced cell area and AR, leading to decreased vessel branching density (A-26.83, G-16.66) in both regions on d10. Conversely, non-treated high-resolution designs displayed vessel remodeling with increased CD31+ cell area (1384 µm^2^) and AR (3.00) on d10, facilitating vessel regression into thick singular vessels elongated along the printing direction with minimal branching density (A-43.96, G-31.94) in aptamer regions.

At medium-resolution, CD31+ cells within aptamer regions exhibited higher cell areas (2412 µm^2^) and AR (3.54) in CS-treated samples compared to non-treated samples (area-2296 µm^2^; AR-2.96) on d5. The high CD31+ cell area with elongated/sprouting morphology in CS-treated samples led to thin vessel network structures with large intervessel spaces, low vessel densities (A-22.16%; G-5.38%), and branching densities (A-244.3; G-11.06) in the aptamer regions on d5. In contrast, the lower cell area and elongation in non-treated samples resulted in densely packed vessel networks with minimal intervessel spaces, higher vessel densities (A-27.97%; G-6.28%), and branching densities (A-328.3; G-22.80) on d5. Additionally, this difference in vessel arrangement led to longer average vessel lengths in non-treated samples (A-0.221mm; G-0.028mm) compared to CS-treated samples (A-0.149mm; G-0.022mm) on d5. Over time, the CD31+ cell areas (2575 µm^2^) increased with a statistically insignificant decrease in AR (3.33) in CS-treated samples, while non-treated samples showed increased cell areas (2513

µm^2^) and AR (3.09) by d10. These changes in cell properties became more pronounced as the network organization drastically evolved over time (Fig. 7A&S4). The initial vascular plexus with large intervessel spaces in CS-treated samples remodeled into a honeycomb-shaped network, where individual EC networks joined to form hierarchically spaced intervessel spaces by d10. Conversely, no significant vessel network remodeling was observed in non-treated samples, which exhibited densely packed vessels that regressed near the interface region by d10 (Fig. S4). These morphological changes were further supported by significantly increased average vessel lengths [+CS(A-0.536mm, G-0.114mm); –CS(A-0.167mm, G-0.031mm)] and branching densities [+CS(A-266.9, G-97.41); –CS(A-192.2, G-20.56)] in CS-treated samples compared to non-treated samples on d10.

Following the same trend, low-resolution designs demonstrated a clear advantage of CS-treatment in guiding vessel network organization over time. On d5, CD31+ cells in low-resolution samples displayed almost similar cell areas (+CS: 2497 µm^2^; –CS: 2447 µm^2^) and AR (+CS: 3.54; –CS: 3.55), irrespective of CS-treatment. However, in CS-treated samples, these cells formed highly dense vessel networks with a clear hierarchy of vessel branching, where thin vessel networks merged into thicker branches on d5. This effect was especially prominent near the interface region (Fig. 6&S4, white arrows). In contrast, non-treated samples formed densely packed networks with less prominent CD31+ staining compared to medium-resolution designs, suggesting vessel breakage and regression on d5. Low vessel reorganization and increased regression near the interface region were observed in low-resolution, non-treated samples compared to medium-resolution designs. Over-time, in CS-treated samples, the network remodeled into a hierarchically arranged self-organized structure with varying intervessel spaces, where thin EC networks merged into thick vessel branches by d10. Interestingly, CD31+ cell characteristics remained similar in CS-treated samples on d10, whereas non-treated samples exhibited a drastic decrease in cell area and an increase in AR on d10. These changes in cell properties could be attributed to the overall lower vessel lengths and branching density in non-treated samples on d10. Vessels in non-treated, low-resolution samples reorganized into thicker networks with low branching density and high intervessel spaces by d10. For instance, CS-treated low-resolution designs showed significantly higher branching density [+CS(A-428.9, G-16.53); –CS(A-143.1, G-11.14)] and average vessel lengths [+CS(A-0.774mm, G-0.076mm); –CS(A-0.204mm, G-0.062mm)] compared to non-treated samples on d10.

During embryonic development, EC coalesce to form a honeycomb-shaped primitive capillary plexus through vasculogenesis.[52] Over time, this primary plexus undergoes vessel regression or pruning of immature vessels in the inner plexus, leading to increased intervessel spaces around major vessels. Vessel remodeling is crucial for shaping hierarchically organized, multiscale vascular networks *in vivo*. Remarkably, the aptamer regions of the CS-treated medium and low-resolution designs exhibited similar behavior *in vitro*. Specifically, in low-resolution designs, the vascular plexus observed on d5 underwent substantial network remodeling by d10, resulting in increased vessel density (1.35-fold), branching density (1.54-fold), average vessel length (2.19-fold), and intervessel spaces. This *in vitro* network remodeling was confined to the aptamer regions of the bioprinted constructs and could be temporally modulated with time-controlled CS-treatment.

We postulate that VEGF gradients, facilitated by a combination of CS-triggered released VEGF and aptamer-tethered VEGF in CS-treated samples of medium and low-resolution designs, contribute to higher cell area (low-resolution, d10: 2437 µm^2^) and a relatively curved/elongated morphology of CD31+ cells. This gradient aids in the effective rearrangement of vessels in the aptamer regions over time. In contrast, non-treated samples at both resolutions, lacking this VEGF gradient, exhibited elongated CD31+ cells with lower cell areas (low-resolution, d10: 1049 µm^2^, p<0.001), resulting in densely packed vessel networks with regression and breakage by d10. This suggests that the CS-treatment effect on network organization in all designs is transient, leading to hierarchical vascular network rearrangement, especially in medium and low-resolution designs. Conversely, the prolonged presence of high concentrations of aptamer-tethered VEGF tends to regress these vascular networks, but increasing spatial printing resolution can mitigate vessel regression.

Regarding vessel organization within printed GelMA regions, a common behavior was observed in medium and low-resolution designs. CS-treatment led to minimal network formation on d5, but over time, networks invaded the GelMA regions from the aptamer side via interface. In contrast, non-treated samples showed considerable network formation on d5, which regressed over time. In all cases, thin vessel networks with large intervessel spaces invaded the GelMA regions via the interface. Interestingly, the localization of aptamer-tethered VEGF within bioprinted constructs restricted hierarchical vessel formation to the printed aptamer regions at all spatial resolutions. Thus, programmable bioinks can influence cell behavior over a significant spatial range in GelMA regions (501 to 2223 µm), provided similar spatial ranges of programmable bioinks are present (338 to 2014 µm). Notably, at all resolutions, non-treated samples exhibited enhanced directionality of self-organized networks parallel to the printing direction until d10, though the mechanism behind this directional behavior remains unknown.

CS-triggering guides CD31+ EC orientation and network organization in a time-controlled manner, depending on the programmable bioink’s spatial resolution (Fig. S5). On d5, in CS-treated aptamer regions, high-resolution designs showed CD31+ cells aligned in all directions, medium-resolution designs promoted parallel alignment followed by perpendicular alignment, and low-resolution designs favored maximal perpendicular alignment (Fig. S5B,D&F). In GelMA regions, CD31+ cells predominantly aligned parallelly across all designs on d5. Conversely, non-treated samples displayed higher percentages of perpendicularly aligned cells in the aptamer regions, while GelMA regions had mostly parallelly aligned cells, with more pronounced differences in low-resolution designs. By d10, high-resolution designs showed the maximum number of parallelly aligned cells in both regions, medium-resolution designs had similar amounts of parallel and perpendicular alignment, and low-resolution designs exhibited maximal perpendicular cell alignment. Non-treated high and medium-resolution designs showed enhanced parallel alignment of cells in both regions on d10, with lower cell percentages aligned in other directions. However, non-treated low-resolution designs exhibited a sharp decrease in cell numbers aligned in all direction in the aptamer regions compared to d5, with the majority of cells in GelMA regions aligned parallelly [-CS low-resolution, d10: 0°-20°(A-12.21%, G-22.38%), 160°-180°(A-12.21%, G-13.98%), 80°-100°(A-8.01%, G-11.88%)] (Fig. S5F).

This CD31+ cell alignment behavior in medium and low-resolution designs contributes to vessel network rearrangement and hierarchical organization within aptamer regions by d10. Notably, CS-triggering affects cell orientation in both printed aptamer regions and neighboring GelMA regions. Although the immediate effect of CS-triggering on cell orientation appears transient, long-term analysis reveals that CS-treatment enhances these effects over time, maintain consistent cell alignment trends. CS-treated low-resolution designs facilitated maximal perpendicular cells alignment on d10. Compared to traditional bioinks, where only spatial resolution influences cell orientation through mechanical cues, programmable bioinks use both mechanical and biochemical (aptamer-tethered VEGF) guiding cues synergistically to direct EC network morphogenesis spatiotemporally.

### 2.6 *Neighboring GelMA region and CS synergistically enhance vascular morphogenesis in* bioprinted aptamer region

Next, we investigated how neighboring GelMA regions and design architecture influence vascular morphogenesis within aptamer regions of bioprinted constructs, particularly when treated with CS. We bioprinted vascular tree designs with angles ranging from 40° to 80° (Fig. 8A), reflecting physiological artery bifurcations *in vivo*.[53,54] We categorized filament widths into high (<500 µm), medium (700-1200 µm), and low (>1140 µm) spatial resolutions, similar to previous parallel line designs (Fig. 8B). The printed filament widths averaged 442.9±106.1 µm (CV=23.95%) for high-resolution, 796.8±142.6 µm (CV=17.89%) for medium-resolution, and 1158±134.5 µm (CV= 11.62%) for low-resolution. Using programmable bioink premixed with MSCs/HUVECs, we bioprinted these designs, followed by VEGF loading and CS-triggering on d4. Confocal microscopy revealed positive staining for CD31, F-actin, and Hoechst (nuclei) in all designs, with MSCs maintaining a round morphology irrespective of CS treatment, unlike parallel line designs.

**Figure 8.**
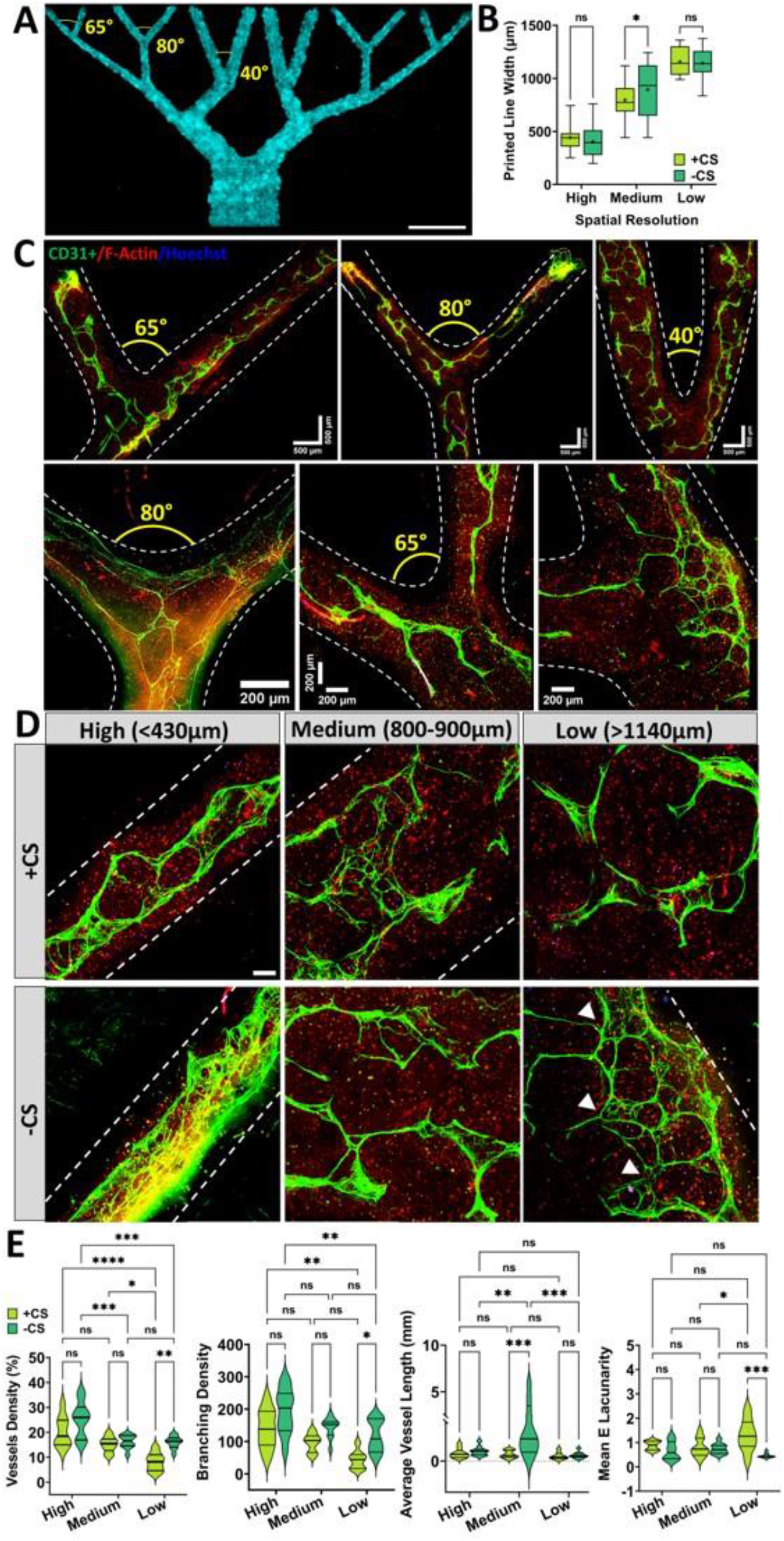
3D-bioprinted vascular tree design using programmable bioink. (A) Stitched fluorescent image of 3D-bioprinted vascular tree design using programmable bioink. The scale bar is 10mm. For comparative analysis, the design was categorized into three distinct regions depending on their printed line widths, i.e., high (<430 µm), medium (800-900 µm) and low-resolution (>1140 µm). (B) Printed line width quantification using two individual bioprinted samples, which were then used for +/-CS studies. (C) Confocal z-stacks stitched mosaics of design without CS treatment on d10. The scale bars are marked in the individual mosaics. The designs were immunostained for CD31 expression (green), cell cytoskeletal F-actin (red) and nuclei (blue). (D) Confocal z-stacks showing different spatial resolution regions among designs treated with/without CS (added on d4) on d10. The scale bar is 100 µm. (E) Vascular network properties as a function of spatial resolution among CS treated/ non-treated samples on d10 (n=6, technical replicates).

EC can detect curvature variations, especially near vessel bifurcations.[55] Confocal microscopy mosaics showed that bioprinted bifurcations serve as templates for developing CD31+ EC networks (Fig. 8C). Different bifurcation angles resulted in distinct network formations, with 65° angles demonstrating optimal network stability. Higher bifurcation angles led to interconnected networks with minimal breakage, whereas lower angles increased network stability.

At bifurcation angles near 80° (with spatial resolution <430 µm), EC networks primarily aligned along the periphery of the printed line, creating large intervessel spaces. At 65° (with spatial resolution <430 µm), EC networks thickened with fewer branches, mainly in the middle of the printed line. At 40° (with spatial resolution 800-900 µm), EC networks formed along the periphery, with interconnections spanning the width of the printed line, but with increased breakages near bifurcations. Higher bifurcation angles thus promoted more stable interconnected networks, while lower angles induced instability and breakage. CS-treatment increased network breakage near 65° bifurcation regions compared to non-treated samples (Fig. S6). This suggests that localized aptamer-tethered VEGF enhances network stability, especially near 65° bifurcations.

EC generate traction forces to mediate cell-cell connectivity and network self-assembly by directing cell migration.[56] Stress fibers, myosins, and focal adhesions facilitate traction force generation. Cells cultured near bifurcations exert lower traction forces compared to those distant from bifurcations.[57] Heterogenous traction force distribution among neighboring ECs creates unstable regions with high force fluctuations, potentially causing inter-endothelial gaps.[58] The increased vascular network breakage near bifurcations may result from asymmetrical traction force distribution and other intracellular forces. Drawing definitive conclusions necessitates further experiments related to traction force measurements.

Similar to parallel line designs, we observed a negative correlation between spatial resolution and network formation in vascular tree designs. Non-treated designs in low-resolutions regions exhibited enhanced network self-assembly with hierarchical organization and intervessel spaces. Near low-resolution bifurcation regions within a single bioprinted line, augmented vessel networks formed along the periphery, with new networks sprouting towards the midsection near bifurcations (Fig. 8C, white arrow). We speculate that this directional sprouting arises from local gradients in aptamer-tethered VEGF concentrations, with high VEGF zones near the bifurcation point and low VEGF zones near the periphery. These results confirm the bioavailability of aptamer-tethered VEGF within programmable bioinks.

Comparing vascular tree and parallel line designs, we focused on CD31+ network organization within the linear regions of vascular tree designs, excluding curvatures. High spatial resolution correlated with enhanced parallel alignment of vascular networks (Fig. 8D). By d10, CS-treated samples showed CD31+ networks with larger intervessel spaces and reduced honeycomb-shaped arrangements compared to non-treated groups. CS-treatment did not significantly affect vessel properties at high spatial resolution by d10, unlike in parallel line designs. At low-resolution, CS-treatment reduced vessel density (+CS: 8.141±3.664%; –CS: 16.03±2.451%; p<0.01) and branching density (+CS: 43.36±28.29%; –CS: 117.1±49.68%; p<0.05) with negligible changes in average vessel length. At medium-resolution, CS-treatment significantly decreased average vessel length (+CS: 0.426±0.24 mm; –CS: 2.087±1.857 mm; p<0.001) with insignificant changes in network branching (Fig. 8E).

Intrigued by the contrasting vessel organization trends, we compared vessel properties on d10 between parallel line (with, w/ GelMA) and vascular tree designs (without, w/o GelMA) in their respective aptamer regions (Fig. S7). High-resolution designs w/o GelMA exhibited significantly higher vessel density than w/ GelMA designs, under both CS-treated (w/ GelMA-6.252±1.13%, CV=18.07%; w/o GelMA-20.24±6.252%, CV=30.89%; p<0.0001) and non-treated conditions (w/ GelMA-7.533±2.36%, CV=31.41%; w/o GelMA-25.36±7.11%, CV=28.02%; p<0.0001).

GelMA regions effectively modulated vessel properties in aptamer regions at medium and low-resolutions. CS-treated samples w/ GelMA showed linear increases in vessel density and branching density at both resolutions compared to the w/o GelMA group [branching density, (low-resolution: w/ GelMA-428.9; w/o GelMA-43.36; p<0.0001)(medium-resolution: w/ GelMA-266.9; w/o GelMA-96.38; p<0.0001)]. These CS-treated w/ GelMA designs had significantly shorteraverage vessel lengths than w/o GelMA designs. Thus, at medium and low-resolutions, the presence of GelMA regions combined with CS-treatment enhanced vessel network and branching while maintaining shorter average vessel lengths, leading to hierarchical network arrangements. Non-treated w/ GelMA designs at both resolutions showed slightly higher vessel properties, including average vessel length, compared to w/o GelMA designs, though these differences were statistically insignificant. GelMA regions enhanced network organization in aptamer regions only at medium and low-resolutions, with no influence at high-resolution (<500 µm). CS-treatment combined with GelMA regions at medium and low-resolutions effectively organized vessel networks with hierarchical branching and density. The presence of GelMA regions at these resolutions facilitated increased vessel properties, although differences were insignificant without CS-treatment, highlighting the importance of CS-mediated VEGF release.

The shape and stability of VEGF gradients critically influence EC proliferation and migration. Computational models suggest that VEGF distribution forms an even slope along the gradient, with cells experiencing exponentially higher concentrations near the source and linearly lower concentrations further away.[59] EC migration in response to VEGF is more pronounced in exponential regions than in linear regions, with chemotaxis diminishing as cells reach the high end of the gradient. Cells can also sense changes in gradient steepness, reducing net migration as they approach higher VEGF concentrations.[59] Programmable bioink designs with varying spatial resolutions generate VEGF gradients of different steepness, especially in CS-treated designs. Low-resolution designs result in steeper concentration gradients (Fig. 3), forming stable, hierarchically organized vascular networks within aptamer regions (Fig. 7). We propose that neighboring GelMA regions act as buffering zones, facilitating stable VEGF gradient formation within bioprinted constructs. The lack of organized vascular networks in designs without GelMA regions highlights the importance of these buffering zones. The ability of bioprinted designs to maintain stable VEGF gradients up to six days after CS-treatment contributes to vascular network remodeling, stabilization and maturation. In tissue microenvironments, a combination of spatiotemporally controlled chemotactic and haptotactic cues guides EC morphogenesis.[60] Similarly, in our bioprinted constructs, aptamer-tethered VEGF molecules within the ECM (haptotactic cue) and freely diffusing VEGF molecules upon CS-triggering (chemotactic cue) synergistically form stable concentration gradients and guide network morphogenesis. Without CS-triggering, vascular networks show reduced branching and higher alignment within the constructs, emphasizing the importance of freely diffusing chemotactic cues in guiding vascular morphogenesis and establishes CS an a temporally controlled external trigger.

Matching the vascular network organization and orientation with the host tissue promotes the anastomosis of engineered networks *in vivo*.[3,61] In this study, we demonstrated that using programmable bioinks in combination with buffering GelMA regions and CS-triggering can effectively modulate vascular network orientation and organization transiently within spatially localized regions of engineered tissues, in a time-controlled manner. To elicit these transient and selective changes, we employed external triggers. These programmable bioinks with external trigger technology, unlike conventional bioinks, not only allow for spatial control but also provide temporal control over the in-situ manipulation of engineered vasculature days after implantation. Programmable bioinks offer significant advantages over current strategies for controlling vascular network morphogenesis within the actively remodeling microenvironment of implanted engineered tissue. We believe that CS-triggered control over network orientation and organization can be used to match host tissue vasculature, thereby facilitating anastomosis *in vivo*.

We can design aptamer sequences to capture and release various GFs with ease, making programmable bioinks highly adaptable for any GF of choice. Owing to the aptamer’s high binding specificities, we can incorporate multiple aptamers within the same bioink, each engineered for different release kinetics for various GFs. In future studies, we will combine multiple GFs using these bioinks to better regulate the multiple stages of vascular morphogenesis and anastomosis.

Furthermore, conventional approaches often necessitate the use of supraphysiological GF concentrations, which can be detrimental *in vivo*, leading to construct failure and regressed anastomosis. Physiological levels of VEGFA in healthy humans range from 1-3 pM in blood plasma and tissue interstitial fluid, increasing in response to hypoxia.[8] State-of-the-art bioprinted constructs use supraphysiological concentrations ranging from 25 ng up to 3 µg of VEGF for *in vitro* and *in vivo* analysis.[18,19,62] However, our work employed very low VEGF concentrations, significantly reducing the risk of off-target effects associated with GF delivery. This approach not only enhances the precision of vascular network formation but also underscores the efficiency and safety of our programmable bioinks in tissue engineering applications.

## 3. Conclusions

We developed a programmable bioink that dynamically present VEGF to guide vascular morphogenesis within bioprinted constructs in a spatiotemporally controlled manner. This approach leverages the aptamer’s high affinity to rapidly sequester VEGF in a specific region of the construct and utilizes aptamer-CS hybridization to control the release of VEGF from that region in a time-controlled manner, even days after bioprinting. Although aptamers-mediated VEGF delivery in hydrogels and its effect on EC network formation have been studied previously, our data are the first, to our knowledge, to demonstrate how changes in spatial resolution of programmable bioink within a bioprinted construct, combined with CS-triggered VEGF release, significantly affects the alignment, organization, and morphogenesis of microvascular networks in a time-controlled manner. The presence of aptamer-tethered VEGF and the generation of instantaneous VEGF gradients upon CS-triggering restrict hierarchical network formation to the printed aptamer regions at all spatial resolutions, demonstrating the capability of programmable bioinks to influence network properties over significant spatial ranges. Network properties improve as the spatial resolution of the programmable bioink decreases, with low-resolution designs showing the highest network properties. Specifically, low-resolution designs subjected to CS-treatment exhibit substantial vascular network remodeling, characterized by an increase in vessel density (1.35-fold), branching density (1.54-fold), and average vessel length (2.19-fold) compared to non-treated samples, where these properties remained unchanged. This indicates that CS acts as an external trigger that can induce transient changes in network organization on-demand, within spatially localized regions of a bioprinted construct. We envision that programmable bioinks will offer new opportunities to bioengineer functional vascular networks within engineered tissues, advancing the field of complex tissue engineering.

## 4. Materials and methods

For a complete list of materials and experimental detail of the present study, please refer to the S1 section of Supplementary Information file that is available online.

## CRediT author statement

**Deepti Rana:** Conceptualization, Methodology, Investigation, Formal analysis, Writing-original draft, Writing – review & editing. **Vincent R. Rangel**: Methodology, Investigation. **Prasanna Padmanaban:** Methodology, Investigation. **Vasileios D. Trikalitis:** Methodology, Investigation. **Ajoy Kandar:** Methodology, Investigation. **Hae-Won Kim:** Writing-review & editing. **Jeroen Rouwkema:** Conceptualization, Writing-review & editing, Supervision, Project administration, Funding acquisition.

## Declaration of competing interest

The authors declare that they have no known competing financial interests or personal relationships that could have appeared to influence the work reported in this paper.

## Data availability

Data will be made available on request.

## Acknowledgements

This work was supported by the European Research Council (ERC) under the European Union’s Horizon 2020 Research and Innovation Program (No. 724469). Illustrations for the manuscript were created with BioRender.com.

## Supplementary Information

Supplementary data to this article can be found online.

## Notes

### Competing Interest Statement

The authors have declared no competing interest.

## References

[1] J. Rouwkema, A. Khademhosseini, Trends Biotechnol 2016, 34, 733.

[2] E. A. Margolis, N. E. Friend, M. W. Rolle, E. Alsberg, A. J. Putnam, Trends Biotechnol 2023, 41, 1400.

[3] T. Mirabella, J. W. Macarthur, D. Cheng, C. K. Ozaki, Y. J. Woo, M. T. Yang, C. S. Chen, Nature Biomedical Engineering 2017 1:6 2017, 1, 1.

[4] L. Debbi, B. Zohar, M. Shuhmaher, Y. Shandalov, I. Goldfracht, S. Levenberg, Biomaterials 2022, 280, 121286.

[5] A. A. Szklanny, M. Machour, I. Redenski, V. Chochola, I. Goldfracht, B. Kaplan, M. Epshtein, H. Simaan Yameen, U. Merdler, A. Feinberg, et al., Advanced Materials 2021, 33, 2102661.

[6] S. Ben-Shaul, S. Landau, U. Merdler, S. Levenberg, Proc Natl Acad Sci U S A 2019, 116, 2955.

[7] R. K. Jain, P. Au, J. Tam, D. G. Duda, D. Fukumura, Nature Biotechnology 2005 23:7 2005, 23, 821.

[8] P. S. Briquez, L. E. Clegg, M. M. Martino, F. Mac Gabhann, J. A. Hubbell, Nature Reviews Materials 2016 1:1 2016, 1, 1.

[9] C. Ruhrberg, H. Gerhardt, M. Golding, R. Watson, S. Ioannidou, H. Fujisawa, C. Betsholtz, D. T. Shima, Genes Dev 2002, 16, 2684.

[10] S. P. Herbert, D. Y. R. Stainier, Nat Rev Mol Cell Biol 2011, 12, 551.

[11] L. Pérez-Gutiérrez, N. Ferrara, Nature Reviews Molecular Cell Biology 2023 24:11 2023, 24, 816.

[12] J. Bliley, J. Tashman, M. Stang, B. Coffin, D. Shiwarski, A. Lee, T. Hinton, A. Feinberg, Biofabrication 2022, 14, DOI 10.1088/1758-5090/AC58BE.

[13] V. D. Trikalitis, N. J. J. Kroese, M. Kaya, C. Cofiño-Fabres, S. ten Den, I. S. M. Khalil, S. Misra, B. F. J. M. Koopman, R. Passier, V. Schwach, et al., Biofabrication 2022, 15, DOI 10.1088/1758-5090/ACA124.

[14] F. L. C Morgan, L. Moroni, M. B. Baker, F. L. C Morgan, L. Moroni, M. B. Baker, Adv Healthc Mater 2020, 9, 1901798.

[15] P. Tournier, G. Saint-Pé, N. Lagneau, F. Loll, B. Halgand, A. Tessier, J. Guicheux, C. Le Visage, V. Delplace, Advanced Science 2023, 10, 2300055.

[16] S. M. Hull, J. Lou, C. D. Lindsay, R. S. Navarro, B. Cai, L. G. Brunel, A. D. Westerfield, Y. Xia, S. C. Heilshorn, Sci Adv 2023, 9, DOI 10.1126/SCIADV.ADE7880/SUPPL_FILE/SCIADV.ADE7880_SM.PDF.

[17] A. Mir, E. Lee, W. Shih, S. Koljaka, A. Wang, C. Jorgensen, R. Hurr, A. Dave, K. Sudheendra, N. Hibino, Bioengineering 2023, Vol. 10, Page 606 2023, 10, 606.

[18] F. E. Freeman, P. Pitacco, L. H. A. van Dommelen, J. Nulty, D. C. Browe, J. Y. Shin, E. Alsberg, D. J. Kelly, Sci Adv 2020, 6, 5093.

[19] A. Poerio, J. F. Mano, F. Cleymand, ACS Biomater Sci Eng 2023, 9, 6531.

[20] G. L. Koons, A. G. Mikos, J Control Release 2019, 295, 50.

[21] H. Cui, W. Zhu, B. Holmes, L. Grace Zhang, H. Cui, W. Zhu, B. Holmes, L. G. Zhang, Advanced Science 2016, 3, 1600058.

[22] M. Aliabouzar, A. W. Y. Ley, S. Meurs, A. J. Putnam, B. M. Baker, O. D. Kripfgans, J. B. Fowlkes, M. L. Fabiilli, Bioprinting 2022, 25, DOI 10.1016/J.BPRINT.2021.E00188.

[23] M. Li, Z. Liu, Z. Shen, L. Han, J. Wang, S. Sang, Int J Biol Macromol 2024, 262, DOI 10.1016/J.IJBIOMAC.2024.130075.

[24] M. Falandt, P. N. Bernal, O. Dudaryeva, S. Florczak, G. Größbacher, M. Schweiger, A. Longoni, C. Greant, M. Assunção, O. Nijssen, et al., Adv Mater Technol 2023, 8, DOI 10.1002/ADMT.202300026.

[25] D. Rana, A. Kandar, N. Salehi-Nik, I. Inci, B. Koopman, J. Rouwkema, Bioact Mater 2022, 12, 71.

[26] D. Rana, P. Padmanaban, M. Becker, F. Stein, J. Leijten, B. Koopman, J. Rouwkema, Mater Today Bio 2023, 19, 100551.

[27] C. Liu, Q. Yu, Z. Yuan, Q. Guo, X. Liao, F. Han, T. Feng, G. Liu, R. Zhao, Z. Zhu, et al., Bioact Mater 2023, 25, 445.

[28] S. Asim, T. A. Tabish, U. Liaqat, I. T. Ozbolat, M. Rizwan, Adv Healthc Mater 2023, 12, 2203148.

[29] J. Yin, M. Yan, Y. Wang, J. Fu, H. Suo, ACS Appl Mater Interfaces 2018, 10, 6849.

[30] W. Liu, M. A. Heinrich, Y. Zhou, A. Akpek, N. Hu, X. Liu, X. Guan, Z. Zhong, X. Jin, A. Khademhosseini, et al., Adv Healthc Mater 2017, 6, DOI 10.1002/ADHM.201601451.

[31] F. Zhou, P. Wang, J. Chen, Z. Zhu, Y. Li, S. Wang, S. Wu, Y. Sima, T. Fu, W. Tan, et al., Nucleic Acids Res 2022, 50, 9039.

[32] P. Shi, X. Wang, B. Davis, J. Coyne, C. Dong, J. Reynolds, Y. Wang, Angewandte Chemie International Edition 2020, 59, 11892.

[33] N. K. Singh, Y. Wang, C. Wen, B. Davis, X. Wang, K. Lee, Y. Wang, Nature Biotechnology 2023 2023, 1.

[34] L. Abune, N. Zhao, J. Lai, B. Peterson, S. Szczesny, Y. Wang, ACS Biomater Sci Eng 2019, 5, 2382.

[35] Y. Wang, Biomaterials 2018, 178, 663.

[36] B. G. Soliman, G. S. Major, P. Atienza-Roca, C. A. Murphy, A. Longoni, C. R. Alcala-Orozco, J. Rnjak-Kovacina, D. Gawlitta, T. B. F. Woodfield, K. S. Lim, Adv Healthc Mater 2022, 11, DOI 10.1002/adhm.202101873.

[37] I. Orellano, A. Thomas, A. Herrera, E. Brauer, D. Wulsten, A. Petersen, L. Kloke, G. N. Duda, Adv Funct Mater 2022, 32, 2208325.

[38] E. McEvoy, T. Sneh, E. Moeendarbary, Y. Javanmardi, N. Efimova, C. Yang, G. E. Marino-Bravante, X. Chen, J. Escribano, F. Spill, et al., Nature Communications 2022 13:1 2022, 13, 1.

[39] S. Boularaoui, G. Al Hussein, K. A. Khan, N. Christoforou, C. Stefanini, Bioprinting 2020, 20, e00093.

[40] M. Li, X. Tian, J. A. Kozinski, X. Chen, D. K. Hwang, 10.1142/S0219519415500736 2015, 15, DOI 10.1142/S0219519415500736.

[41] Y. S. J. Li, J. H. Haga, S. Chien, J Biomech 2005, 38, 1949.

[42] M. J. Levesque, R. M. Nerem, Biorheology 1989, 26, 345.

[43] M. Samandari, F. Alipanah, K. Majidzadeh-A, M. M. Alvarez, G. Trujillo-De Santiago, A. Tamayol, Appl Phys Rev 2021, 8, DOI 10.1063/5.0040732.

[44] A. C. Vion, T. Perovic, C. Petit, I. Hollfinger, E. Bartels-Klein, E. Frampton, E. Gordon, L. Claesson-Welsh, H. Gerhardt, Front Physiol 2021, 11, 623769.

[45] U. Nagarajan, G. Beaune, A. Y. W. Lam, D. Gonzalez-Rodriguez, F. M. Winnik, F. Brochard-Wyart, Communications Physics 2021 4:1 2021, 4, 1.

[46] C. Zhang, Z. Zhang, Y. Qi, Polymers 2021, Vol. 13, Page 2605 2021, 13, 2605.

[47] C. Piard, A. Jeyaram, Y. Liu, J. Caccamese, S. M. Jay, Y. Chen, J. Fisher, Biomaterials 2019, 222, 119423.

[48] L. Lamalice, F. Le Boeuf, J. Huot, Circ Res 2007, 100, 782.

[49] J. M. Lee, W. Y. Yeong, J R Soc Interface 2020, 17, DOI 10.1098/RSIF.2020.0294.

[50] H. Aubin, J. W. Nichol, C. B. Hutson, H. Bae, A. L. Sieminski, D. M. Cropek, P. Akhyari, A. Khademhosseini, Biomaterials 2010, 31, 6941.

[51] O. S. Al-Kadi, D. Watson, IEEE Trans Biomed Eng 2008, 55, 1822.

[52] C. Korn, H. G. Augustin, Dev Cell 2015, 34, 5.

[53] Z. Knobel, C. J. Kellenberger, T. Kaiser, M. Albisetti, E. Bergsträsser, E. R. Valsangiacomo Buechel, Int J Cardiovasc Imaging 2011, 27, 385.

[54] S. N. Wright, P. Kochunov, F. Mut, M. Bergamino, K. M. Brown, J. C. Mazziotta, A. W. Toga, J. R. Cebral, G. A. Ascoli, Neuroimage 2013, 82, 170.

[55] M. Luciano, C. Tomba, A. Roux, S. Gabriele, Nature Reviews Physics 2024 2024, In press, 1.

[56] J. P. Califano, C. A. Reinhart-King, J Biomech 2010, 43, 79.

[57] Y. Zhang, N. V. Menon, C. Li, V. Chan, Y. Kang, Biomater Sci 2016, 4, 430.

[58] E. T. Valent, G. P. van Nieuw Amerongen, V. W. M. van Hinsbergh, P. L. Hordijk, Exp Cell Res 2016, 347, 161.

[59] I. Barkefors, S. Le Jan, L. Jakobsson, E. Hejll, G. Carlson, H. Johansson, J. Jarvius, W. P. Jeong, L. J. Noo, J. Kreuger, Journal of Biological Chemistry 2008, 283, 13905.

[60] S. SenGupta, C. A. Parent, J. E. Bear, Nature Reviews Molecular Cell Biology 2021 22:8 2021, 22, 529.

[61] D. Rosenfeld, S. Landau, Y. Shandalov, N. Raindel, A. Freiman, E. Shor, Y. Blinder, H. H. Vandenburgh, D. J. Mooney, S. Levenberg, Proc Natl Acad Sci U S A 2016, 113, 3215.

[62] B. Byambaa, N. Annabi, K. Yue, G. Trujillo-de Santiago, M. M. Alvarez, W. Jia, M. Kazemzadeh-Narbat, S. R. Shin, A. Tamayol, A. Khademhosseini, Adv Healthc Mater 2017, 6, DOI 10.1002/ADHM.201700015.

