## Supplementary Information for "Bioprinting of aptamer-based programmable bioinks to modulate multiscale microvascular morphogenesis in 4D"

#### S1. Materials and methods

##### S1.1. Materials

Type A 300 bloom porcine skin gelatin (G1890-500G, Sigma Aldrich), Methacrylic anhydride (MA, 276685-500ML, Sigma Aldrich), Dulbecco's phosphate buffered saline (DPBS, D8537-500ML, Sigma Aldrich), Fisherbrand™ regenerated cellulose dialysis tubing (12-14 kDa, 21-152-14, Fisher Scientific), bovine serum albumin (BSA, A9418, Sigma Aldrich), tris(2,2'-bipyridyl)dichloro-ruthenium (II) hexahydrate (224758, Sigma), sodium persulfate (S6172, Sigma), VEGF specific 5'acrydite-modified aptamer (47-nt, DNA, IDT), complementary sequence (CS, 46-nt, DNA, IDT), 5'Alexa Fluor 488 modified complementary sequence ($CS_F$, 46-nt, DNA, IDT), nuclease free water (11-04-02-01, IDT), 3-(trimethoxysilyl)propyl methacrylate (TMSPMA, 440159, Sigma-Aldrich), Fluoro-Max dyed blue aqueous fluorescent particles (2µm diameter, B0200, Thermo Scientific), ROKIT INVIVO bioprinter (South Korea), 1 ml Luer-Lock syringe (9210927, BD), 22G standard blunt needles (3914, Cellink), microlance 25G (1506, BD), 24x24mm glass coverslips (Thermo scientific), LED light (Jobmate 20W Lamp), Alexa Fluor® 647 Anti-VEGFA antibody [EP1176Y] (ab206887, abcam), human umbilical vein endothelial cells (HUVECs, C2519A, Lonza), human mesenchymal stromal cells (MSC, PT-2501, Lonza), α –MEM medium (+nucleosides, 22571-020, Gibco), fetal bovine serum (FBS, F7524, Sigma), GlutaMax™ supplement (35050061, Gibco), penicillin-streptomycin (pen/strep, 15140-122, Gibco), L-abscorbic acid (A8960, Sigma Aldrich), trypsin-EDTA 0.25% (+phenol red, 25200072, Gibco), vascular endothelial growth factor 165 human (VEGF, H9166, Sigma Aldrich), Gibco™ FGF-Basic AA 1-155 recombinant human protein (bFGF, PHG0264, Fisher Scientific), endothelial cell basal medium 2 (EGM 2, C-22211, PromoCell), endothelial cell growth medium 2 supplement pack (EGM 2, C-39211, PromoCell), live/dead cell double staining kit (04511, Sigma Aldrich), formaldehyde solution (F8775-25ML, Sigma Aldrich), Triton™ X-100 (T8787, Sigma Aldrich), Invitrogen™ Alexa Fluor™ 647 Phalloidin (A22287, Fisher Scientific), Invitrogen™ hoechst 33342 (H1399, Fisher Scientific), CD31 monoclonal antibody (HEC7, mouse, MA3100, Invitrogen), IgG (H+L) cross-adsorbed goat anti-mouse secondary antibody, Alexa Fluor® 488 (A11001, Invitrogen), Corning® Costar® ultra-low attachment well plates were used for all cell culture experiments (CLS3471 Sigma).

***S1.2 GelMA Synthesis***

Gelatin methacryloyl (GelMA) prepolymer was prepared using a previously described method, employing a medium degree of methacryloyl substitution (~60%).[1–3] In summary, MA (1.25% v/v) was introduced into a gelatin solution (10% w/v) at a flow rate of 0.5 mL/min for one hour at a temperature of 50°C. Subsequently, the reaction mixture was diluted five times using DPBS (40°C), and the resulting solution was subjected to dialysis for a week at 40°C using dialysis tubing with a molecular weight cutoff of 12-14kDa to eliminate any remaining salts. Afterward, the solution was freeze-dried for a week and stored at -20°C.

***S1.3 3D-Bioprinting process***

Three-dimensional (3D) bioprinting experiments were performed using INVIVO Bioprinter (ROKIT Healthcare, South Korea) with two motor-driven extrusion printheads. For bioprinting with two bioinks simultaneously, a dual extrusion setup is required where both extrusion printheads were loaded with 1ml BD Luer-lock syringe fitted with a SLS 3D-printed nylon adaptor, and filled with respective bioinks (programmable and GelMA bioinks). Prior to bioprinting, the bioprinter chamber was thoroughly cleaned with ethanol, disinfected using in-built 45 minutes UV light sterilization cycle and calibrated before each printing cycle. The design G-codes used for operating bioprinter were tailor-made to achieve best print quality using NewCreatorK (version 1.57.68) and Notepad++ (free 3rd party software) with 10mm/s printing speed and 100% infill. The BD Luer-lock syringes were loaded with respective bioinks using Microlance needles with total volume of 1ml. The needle diameter used for 3D-printing all designs was 435 µm. To achieve the desired viscosity of bioinks with lower concentrations of polymer (5% GelMA) suitable for 3D-bioprinting, the loaded syringes were cooled at 4°C for 20 minutes and then immediately used for bioprinting. The printbed temperature was set at 4°C to maintain fidelity throughout printing duration. Due to the presence of photosensitive crosslinkers in with bioinks, the syringes were covered with aluminum foils. To improve the printing efficiency, a custom-made 24x24mm coverslip holder with 9 placeholders was 3D-printed using PLA. Furthermore, the coverslips were treated with TMSPMA as per the protocol described elsewhere,[3] and sterilized using autoclave to ensure better adhesion between the printed construct and the glass surface. The prints were initiated using NewCreatorK software inbuilt with ROKIT bioprinter. To ensure better print fidelity, visible light (400-450 nm, 50mW/cm$^2$) was shone onto the prints throughout the printing process. Afterwards, the prints were shifted to clean laminar hood and subjected to post-print photocrosslinking for 2 minutes using the same visible light. Subsequently, each bioprinted coverslip was placed onto an ultra-low attachment 6-well plates and supplemented with 1ml of medium (or co-culture medium). To investigate the effect of spatial resolution of localized aptamer-tethered VEGF on cell behavior, the following three designs with varying spatial resolutions were printed: high-resolution (one line of programmable bioink was printed adjacent to one line of GelMA bioink; in this pattern total 18 lines were printed), medium-resolution (three lines of programmable bioink was printed adjacently to three lines of GelMA bioink; with this pattern in total 18 lines were printed) and low-resolution (five lines of programmable bioink was printed adjacently to five lines of GelMA bioink; with this pattern in total 25 lines were printed). Each design was printed with 2 stacked layers in z-axis. An additional vascular tree was 3D-bioprinted using programmable bioink to evaluate the efficiency of developed programmable bioinks in printing complex angles/design and its effect on cell behavior.

***S1.4 Cell Culture***

Human mesenchymal stromal cells (MSCs, P2-P5) were cultured in α-MEM medium supplemented with 10% (*v/v*) FBC, 2x$10^{-3}$ M L-glutamine, 0.2x$10^{-3}$ M ascorbic acid, 1% (*v/v*) Pen/Strep and 1 ng/ml bFGF. The human umbilical vein-derived endothelial cells (HUVECs, P3-P6) were cultured using EGM-2 medium with 1% (*v/v*) Pen/Strep. Both cell types were cultured in a humidified atmosphere of 5% $CO_2$ at 37°C with medium change in every two days and passaged till 80% confluence.

***S1.5 Programmable bioink preparation***

For the 3D bioprinting of constructs using two different materials, the bioinks were prepared separately. The programmable bioink was created by mixing GelMA pre-polymer solution (25 µl, 5% w/v GelMA in DPBS) with aptamers (25 µM), and visible light photo-initiators i.e., tris(2,2'-bipyridyl)dichloro-ruthenium(II) hexahydrate, and sodium persulfate (Ru/SPS, 1/10 mM) in an Eppendorf tube to yield a total volume of 50 µl. Similarly, GelMA bioinks were prepared by combining GelMA pre-polymer solution (50 µl, 5% w/v GelMA in DPBS) with fluorescent blue microbeads (1 drop/ml) and Ru/SPS (1/10 mM) solution in another Eppendorf tube. The GelMA bioinks were mixed with fluorescent microbeads for interface identification between the printed regions. The concentrations of Ru/SPS used were previously optimized for GelMA hydrogels.[2,4]

For the co-culture experiments, a co-culture medium was prepared by adding MSCs medium (α-MEM, 10% FBS, 2mM L-glutamine, 0.2 mM ascorbic acid, and 1% Pen/Strep) with HUVECs medium (EGM-2 Basal medium + 1% Pen/Strep + all EGM-2 supplements except for $VEGF_{165}$) in 1:1 ratio. However, the VEGF supplement (C-30260, PromoCell) was excluded from the HUVECs medium. To prepare the cell-laden bioinks, HUVECs (P4) and MSCs (P4) were trypsinized, counted using trypan blue, and re-suspended in a 1:1 ratio to achieve a total seeding density of 2.5x$10^{6}$ cells/ml. The cell suspension was centrifuged at 300g for 3 minutes at 4°C to form a cell pellet. The supernatant was then replaced with the aptamer-based pre-polymer solution (25 µl of 5% GelMA + 25 µM aptamer solution + Ru/SPS 1/10 mM in DPBS) for the programmable bioink and the GelMA-based pre-polymer solution (50 µl of 5% GelMA + blue fluorescent microbeads + Ru/SPS 1/10 mM) for the GelMA bioink. Care was taken while mixing the pre-polymer solutions with the cell pellet to prevent bubble formation. The prepared bioinks were used to fabricate 1-1, 3-3, 5-5, and vascular tree designs, following the procedures described in Section 2.3. After photocrosslinking, the designs were incubated with co-culture medium (1ml) supplemented with VEGF (10ng) at 37°C for 1 hour to allow for growth factor sequestration. The supernatants were then removed, and the designs were washed twice with DPBS before being supplemented with co-culture medium (1ml) without VEGF. The medium was refreshed every 24 hours throughout the study. Furthermore, to investigate the effect of CS-mediated triggered VEGF release on the cell-laden 3D bioprinted designs, 25 µM of CS (aptamer to CS ratio of 1:1) was added to the co-culture medium (1ml) on day 4 of the study.

##### *S1.6 Aptamer-CS molecular recognition post bioprinting*

To assess the aptamer's retention within printed structures and their ability for molecular recognition with CS after the bioprinting process, we employed the 1-1 design. The 1-1 designs were 3D-bioprinted using programmable and GelMA bioinks. Subsequently, these designs were supplemented with Alexa Fluor-488 labeled complementary sequence ($CS_F$) (25µM) in DPBS (1ml) for 24 hr at 37°C. Following this incubation period, the supernatant was discarded and the constructs were washed with DPBS to eliminate any excess or unbound $CS_F$. The washing step was performed using 1 ml of DPBS, and the constructs were imaged using a fluorescence microscope (EVOS M7000, ThermoFisher Scientific) with a 4x objective lens. Two experimental replicates of the 1-1 design were used for comparative analysis. The measure of aptamer-CS molecular recognition was evaluated by quantifying the fluorescence intensity of the sequestered $CS_F$ in the printed regions of both bioinks throughout the samples and as a function of distance using ImageJ software (NIH, USA). Additionally, an additional parameter, print fidelity, was assessed by quantifying and comparing the printed line width of both bioinks among all the designs (1-1, 3-3, and 5-5) using two experimental replicates (n=2).

##### *S1.7 Rheological evaluation*

Programmable bioink (i.e., Aptamer) and GelMA bioink (i.e., GelMA + Bead) were tested for their rheological properties using the stress-controlled rheometer (Physica MCR301, Anton Paar) with parallel plate geometry (PP8, 8mm) at room temperature (293K). The parallel plate and bottom rheometer plate were blasted with sandpaper to prevent wall slip. The plain GelMA bioink without fluorescence microbeads (i.e., GelMA) was prepared as control. To study the changes in viscoelastic behavior of all bioinks during the photocrosslinking process, the bioink's pre-polymer solutions were loaded between the plate by keeping 1 mm gap in a dark room and the rheological measurements were performed in the presence of a visible light lamp ($50mW/cm^2$) to initiate the photocrosslinking process. A custom-made 3D-printed nylon solvent trap was used after gel formation to ensure minimum solvent evaporation. The oscillatory shear measurements at strain ($\gamma$) 0.1% and frequency (f) of 1Hz were performed during the photocrosslinking process (time sweep). The oscillatory shear measurements, both amplitude sweep (at f=1Hz) and frequency sweep (at $\gamma$=0.1%), were carried out to investigate the viscoelastic behavior of the crosslinked bioinks. The storage modulus (G') and loss modulus (G") were determined as the strain (%) changes during oscillatory shear. The storage modulus of bioinks was determined from the linear viscoelastic region of amplitude sweep data.

##### *S1.8 Cell Viability Assay*

Cell viability within 3D-bioprinted 1-1 designs using programmable and GelMA bioinks were evaluated with Live/Dead assay kit as per manufacturer's protocol. In summary, MSCs/HUVECs based cell-laden bioinks were used to 3D-bioprint 1-1 designs, followed by photocrosslinking and VEGF loading for 1hr. The medium was replaced with co-culture medium without VEGF supplement and the designs were incubated for 24hr at 37°C with 5% $CO_2$. Afterwards, the culture medium from the design samples were replaced with 1ml of staining solution [solution A(2 µl) + solution B(1 µl) in 1ml of DPBS] to be incubated for 20min at 37°C and then imaged using fluorescent microscope. The blue color corresponds to the blue fluorescent microbeads present within GelMA bioink. The total number of cells (red and green) and the number of live cells (green) were counted using ImageJ software. The cell viability (%) was quantified as the percentage ratio of the number of live cells by the total number of cells present within one sample. For quantification, three independent design samples (eight images per sample) were used and reported as mean ± standard deviation (SD) for individual samples.

##### *S1.9 In silico Simulation Model of the free VEGF release*

A reaction-diffusion model was employed to forecast the transportation of free VEGF within the 3D-bioprinted designs encompassing both Aptamer and GelMA regions. Specifically, the model was devised to investigate the dispersion of free VEGF molecules from the Aptamer region, where VEGF is abundant, towards the neighboring GelMA regions, characterized by low VEGF concentrations, under the influence of controlled CS triggering (or in its absence). Additionally, the model aimed to analyze the spatiotemporal distribution of free VEGF within both regions as time progressed. The development of the model for analyzing and predicting the diffusion of free VEGF was based on the following assumptions:

1. The system is in equilibrium at the beginning of the simulation (t = 0 h)
2. Free VEGF transports only results from the diffusion
3. Freely diffusing VEGF molecules are the only mobile entities in the system
4. Aptamers are bound onto the hydrogels and tethers free VEGF molecules
5. Upon CS addition, the free VEGF molecules released in the system
6. Higher amount of free VEGF present in the aptamer functionalized hydrogel compared to GelMA hydrogel compartment
7. Free VEGF diffuses from aptamer functionalized region to bulk GelMA regions.
8. Similar to actual sample dimensions, the aptamer and GelMA regions were simulated as rectangles of 1 mm length and 400 µm breadth having placed as aptamer line in middle and GelMA lines on both sides with an interface, in 2D simulations.
9. For 3D simulations, three lines as continues regions placed next to each other in cuboid shape with each region having the following dimensions were used: x=500 µm, y=8 mm & z=800 µm.

The spatiotemporal change in the free VEGF concentration within the hydrogel due to the diffusion over time is given by following equations:

$$\frac{\partial C_{VEGF-aptamer}}{\partial t} = D_{VEGF+CS} \nabla^2 C_{VEGF-aptamer}$$ **[with CS treatment] – (eq. 1)**

$$\frac{\partial C_{VEGF-GelMA}}{\partial t} = D_{VEGF-CS} \nabla^2 C_{VEGF-GelMA}$$ **[without CS treatment] – (eq. 2)**

Where parameters such as $C_{VEGF\text{-}aptamer}$ and $C_{VEGF\text{-}GelMA}$ denotes free VEGF concentration in aptamer and GelMA regions at $t$ = 0 hr, respectively. Additionally, $D_{VEGF+CS}$ and $D_{VEGF\text{-}CS}$ stands for free VEGF diffusivity in the presence and absence of CS throughout the micropattern, respectively **(Table S2, Supplementary Information)**. In both 2D and 3D simulations, we used transport of diluted species physics module in COMSOL (version 5.6) and physics controlled extremely fine mesh conditions was applied to the models for computing free VEGF diffusion.

***S1.10 Immunostainings***

For immunostaining, all samples underwent DPBS rinsing and were subsequently fixed in a 4% formaldehyde solution for 30 mins. After the fixation step, the samples were washed with DPBS and the cell membranes were then permeabilized using 0.1% Triton X-100 for 10 mins. Following this, the samples underwent three subsequent washes with DPBS and were subsequently blocked using a 1% FBS solution for another 45 mins. In the case of actin cytoskeleton staining, the samples were incubated with Alexa Fluor-647 Phalloidin (1:40 dilution in DPBS) for 1 hr at room temperature. This was followed by a 10 mins incubation with Hoechst 33342 (1:2000 dilution in DPBS). In the case of immunostainings, after the blocking step, the samples were washed with DPBS and then incubated overnight at 4°C with a CD31 monoclonal antibody (mouse) (1:200 dilution in DPBS). The samples were subsequently washed with DPBS and supplemented with an Alexa Fluor 488 anti-mouse secondary antibody solution (1:1000 dilution in DPBS) in dark. For samples that underwent double staining with F-actin and CD31, Phalloidin 647 (1:40 dilution in DPBS) was added to the secondary antibody staining solution after 1 hr incubation and allowed to incubate for an additional hour in darkness. Following this, the samples were washed and Hoechst 33342 (1:2000 dilution in DPBS) was added for 10 mins in dark. Once the staining process was completed, the samples were washed, supplemented with 1 ml of DPBS, and stored at 4°C until they were ready for imaging.

***S1.10 Image Analysis***

Image analysis was conducted using both fluorescence microscope image data and maximum projection of confocal z-stacks from all samples, as indicated in the relevant figures, to visualize the filamentous actin and CD31+ cells. ImageJ software was employed for image pre-processing and quantification of cellular properties, including cell area, aspect ratio (AR), and orientation. AR defined as the ratio of the cellular major and minor axis, estimates cell elongation; an AR closer to 1 indicates a circular shape, while >1 value indicates cell stretching/elongation To begin, the single-channel raw image data underwent pre-processing using thresholding, binary, and morphological segmentation tools available in ImageJ. Following image pre-processing, the images were utilized for the analysis of cytoskeletal F-actin stained cells and CD31+ endothelial cells/network properties. The orientation of cells was determined by employing the built-in functions of ImageJ, wherein more than 50 cells (or CD31+ vessel networks) were selected using the wand tool from each segmented image (within both the Aptamer and GelMA regions) to best fit an ellipse with primary and secondary axes. The cell angle (ranging from 0° to 180°) was calculated as the angle between the primary axis and a line parallel to the x-axis of the image. The resulting orientation data was presented as polar plots, with "θ" and "r" representing the cell orientation angle and frequency percentage binned in 30° increments, respectively. Similarly, the pre-processed and segmented images were utilized to quantify CD31+ endothelial vessel network properties, including (the percentage area occupied by CD31+ vessel within total image area, %vessels/total area), average vessel length (mean length of all the vessels in the image), branching density (number of vessel branching points normalized per unit area) and mean lacunarity (nonuniformity or irregular spatial distribution among vascular patterns), using Angiotool software. Vessel density was defined as the percentage of area occupied by CD31+ vessels within the total image area (%vessels/total area), branching density as the number of vessel junctions per area, and average vessel length as the mean length of all vessels in the image. All quantifications were performed with six technical replicates. The obtained data was visualized using GraphPad Prism and Origin software. Additionally, the mosaic tool in ImageJ was employed to stitch together maximum projections of confocal z-stacks for samples depicting the vascular tree design.

##### *S1.11 Statistical Analysis*

The results were expressed as mean ± SD. Statistics was performed using the following variables: (i) for two groups and one time-point, a standard two-tailed, Welch's *t* test was performed, (ii) for more than two groups and one time-point, a one-way analysis of variance (ANOVA) with Dunnett's multiple comparison test was performed, and (iii) when there were more than two groups and multiple time-points, a two-way ANOVA with Tukey's multiple comparisons test was performed. All analyses were performed using GraphPad Prism software. For all comparisons, the alpha was fixed at 0.05 with ***p=<0.0005, ****p=<0.0001 and ns stands for not significant.

### Supplementary Figures

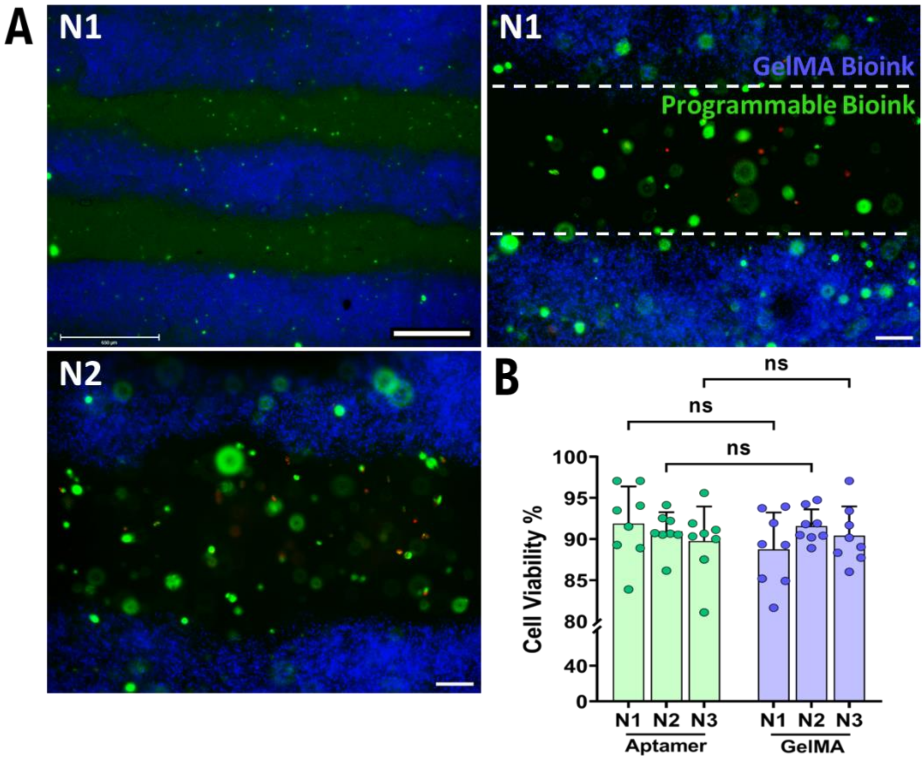


***Figure S1. Cell viability of programmable bioink.*** *Cell viability of the 3D bioprinted construct with HUVECs and MSCs co-cultures (1:1) after 24hr of culture. The bioprinted constructs were visible light crosslinked, followed by VEGF loading for 1hr, washing and subsequently, incubated with co-culture medium for the experiment duration. (A) Live/Dead-stained fluorescent microscopic images of 3D-bioprinted samples using plain GelMA (with blue microbeads) and aptamer-based programmable bioinks at 4x magnification (n1)) and 10x magnification (n1 & n2). The green and red colors represent live and dead cells, respectively. The white dotted line distinguishes the 3D bioprinted GelMA and aptamer regions. The scale bar is 500 µm & 100 µm. (B) Cell viability% quantification was performed using ImageJ software. The cell viability% data GelMA and aptamer regions within three 3D-bioprinted samples are represented as mean±SD, respectively. The statistical significance was calculated using two-way ANOVA with Tukey's multiple comparisons test where alpha was fixed at 0.05 with ***p=0.0005, ****p<0.0001 and ns stands for not significant.*

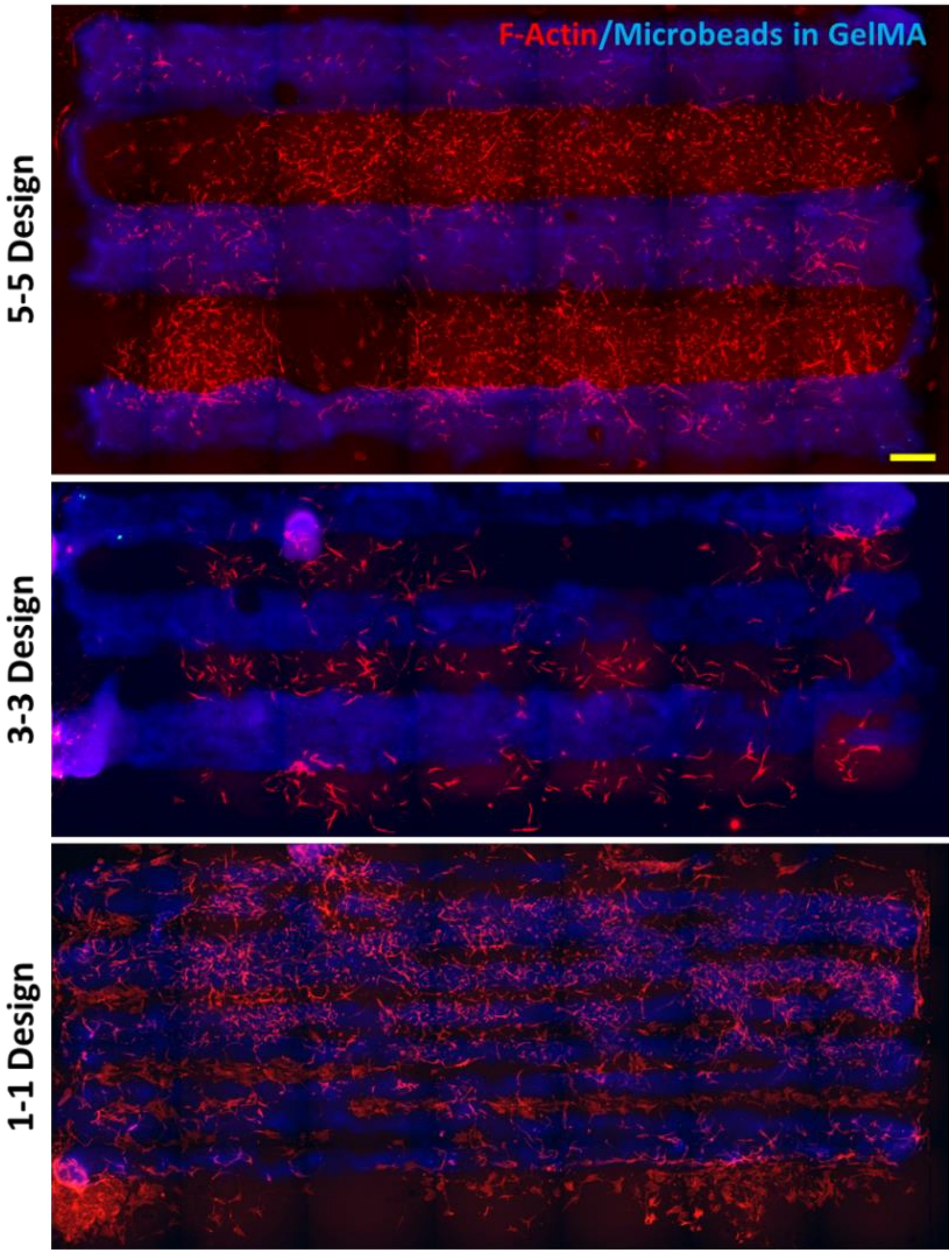


***Figure S2. 3D-Bioprinted designs with MSCs/HUVECs co-culture on D10.*** *The microscopic, stitched images of high-resolution (1-1), medium-resolution (3-3) & low-resolution (5-5) bioprinted designs on D10. The red color corresponds to the cell cytoskeleton F-actin and blue color to the fluorescent microbeads present within GelMA regions of the printed designs.*

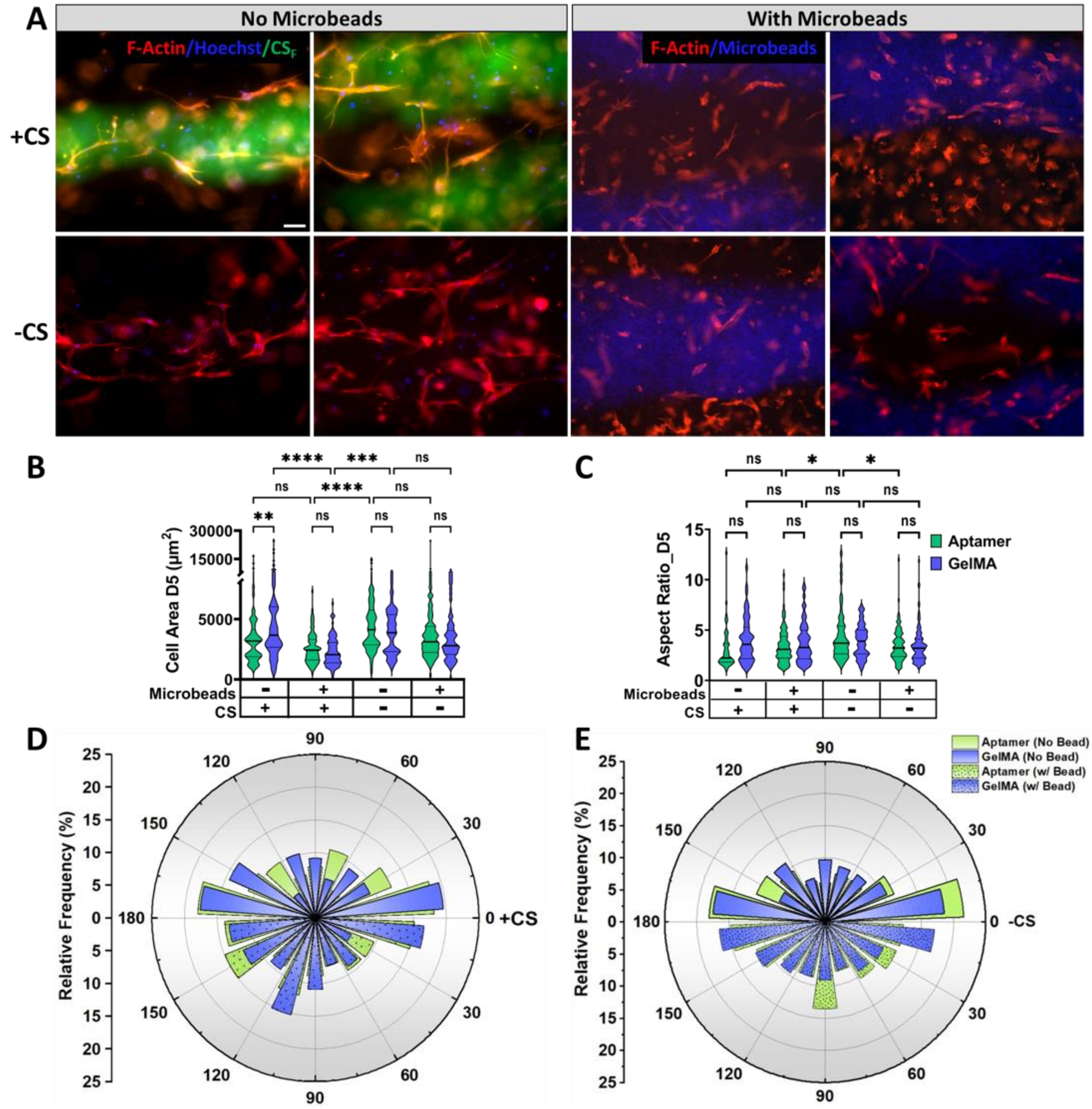


***Figure S3. Comparative analysis of the presence of blue microbeads on the cellular behavior.*** *(A) Representative microscopic images of cell cytoskeleton F-actin (red) within 3D bioprinted constructs in the presence/absence of blue microbeads mixed with GelMA bioink on D5. The designs were supplemented with CS on D4 for triggered VEGF release on D5. The green color corresponds to $CS_F$ used only within no microbead samples for interface visualization among aptamer and GelMA regions. The scale bar is 100 µm. The cell properties such as (B) area, (C) aspect ratio and (D & E) alignment in the presence/absence of CS were evaluated using ImageJ software. The data was calculated using six technical replicates, n=6. The statistical significance was calculated using two-way ANOVA with Tukey's multiple comparisons test where *p<0.05, **p<0.01, ***p<0.001, ****p<0.0001 and ns stands for not significant. The polar plots represent "θ" & "r" as cells alignment and relative frequency % binned in 30º increments in the presence/absence of CS (+/-CS), respectively.*

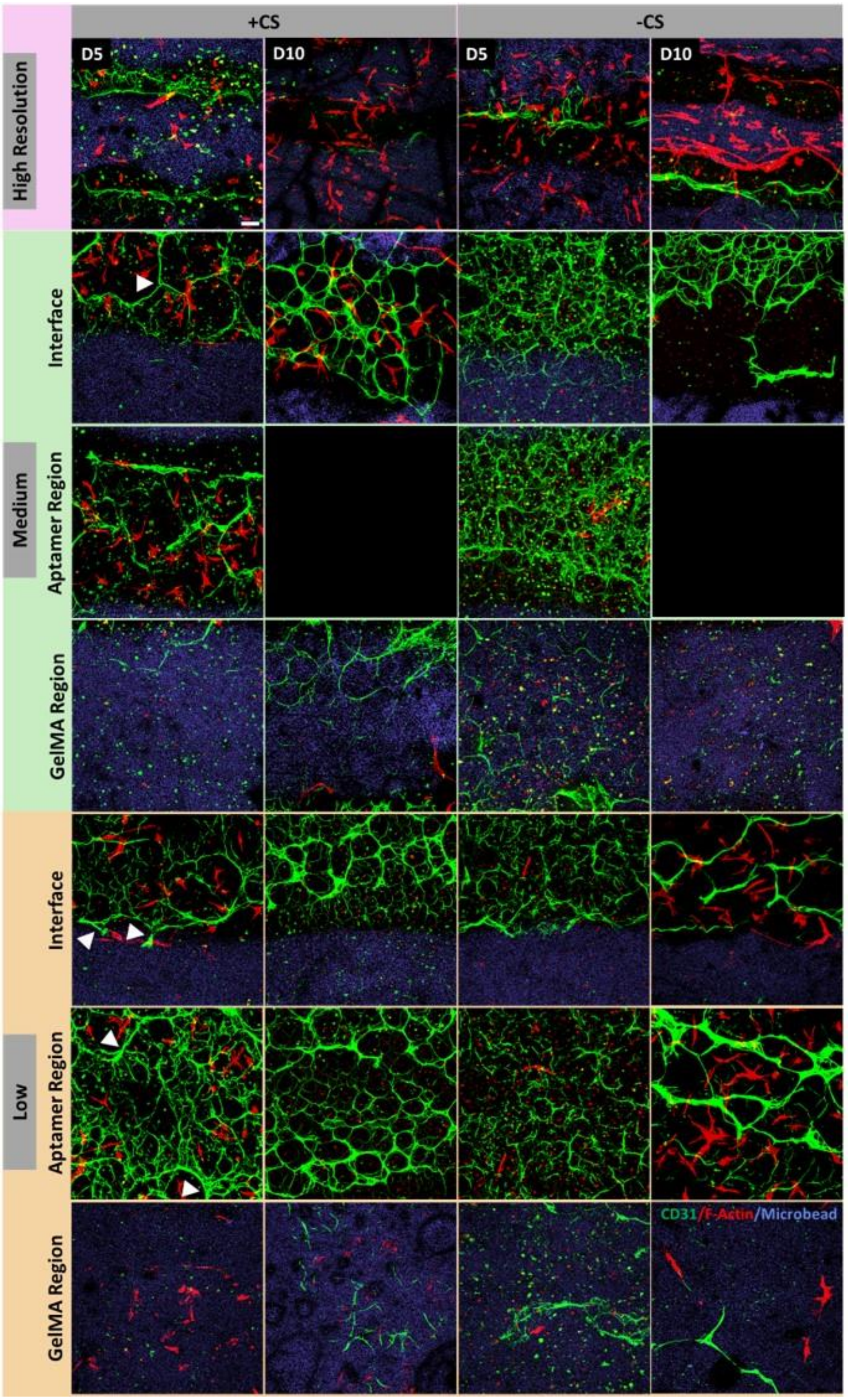


***Figure S4.*** *Maximum projections of confocal z-stack images showing cell cytoskeletal F-actin (red) and endothelial cells specific CD31 (green) expression within MSCs/HUVECs co-cultured, immunostained 3D bioprinted all designs [(high, medium & low-resolution] in the presence/absence of CS (added on D4), on D5 & D10. Each design's interface, aptamer and GelMA regions were shown for comparison. The blue color corresponds to microbeads within GelMA region. The scale bar is 100 µm.*

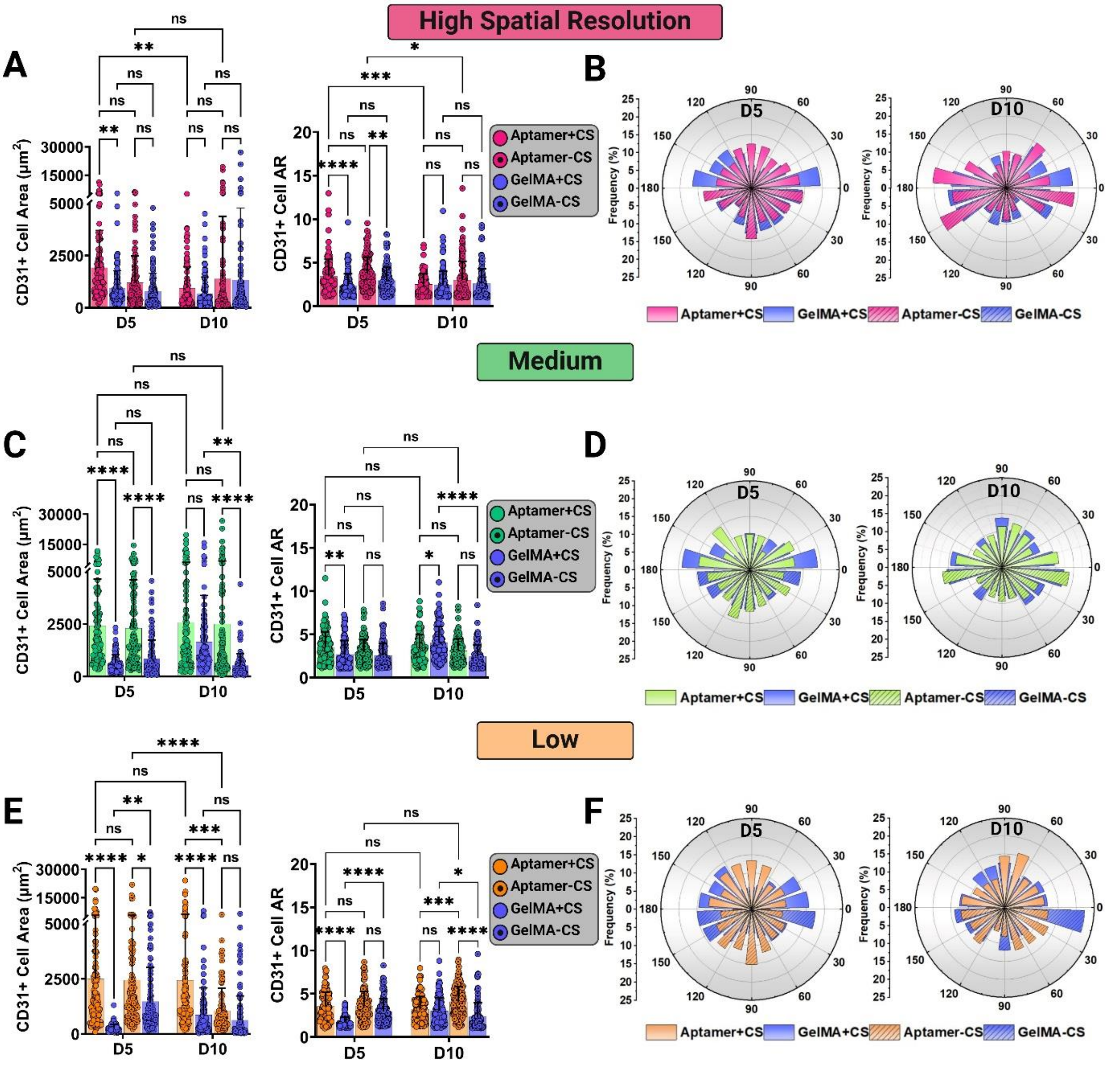


***Figure S5. Comparison of CD31 positive endothelial cell properties among all 3D-bioprinted designs at D5 & D10.*** *CD31+ cell properties such as area, aspect ratio (AR) and alignment on D5 & D10 for (A, B) high-resolution design, (C, D) medium-resolution design and (E, F) low-resolution design, respectively. The cell properties within aptamer and GelMA regions of 3D-bioprinted designs were evaluated using ImageJ with the CD31+ endothelial network shown in confocal microscope images in Figure 6A. The cell area and aspect ratio data are presented as mean±SD with scatter data points. The statistical significance was calculated using two-way ANOVA with Tukey's multiple comparison tests where *p<0.05, **p<0.01, ***p<0.001, ****p<0.0001 and ns stands for not significant. The cell alignment data is represented as polar plot where "θ" & "r" represents cell orientation angle and frequency % binned in 30º increments, respectively. The quantification was performed with six technical replicates, n=6.*

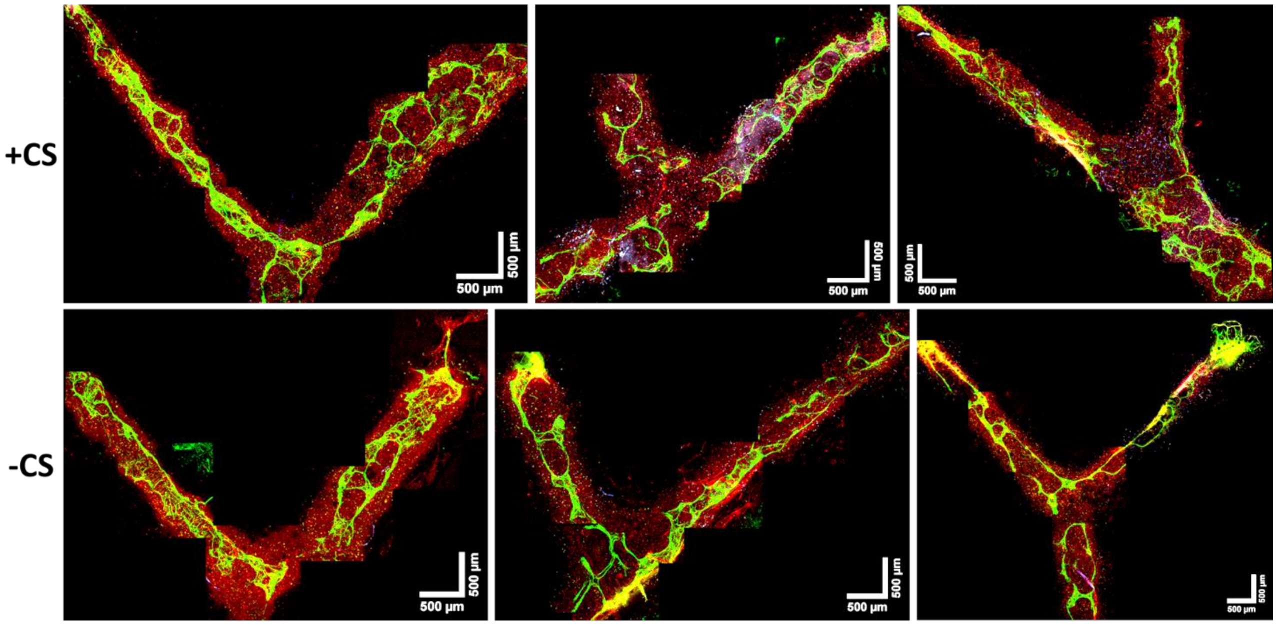


***Figure S6. Stitched mosaics of the maximum projection confocal z-stack images of the vascular tree designs bioprinted using programmable bioink on D10.*** *The figure compares CD31+ endothelial cell (green) organization at the angled bifurcations in the presence/absence of CS (added on D4). The image shows cell cytoskeletal F-actin (red) and cell nuclei (blue).*

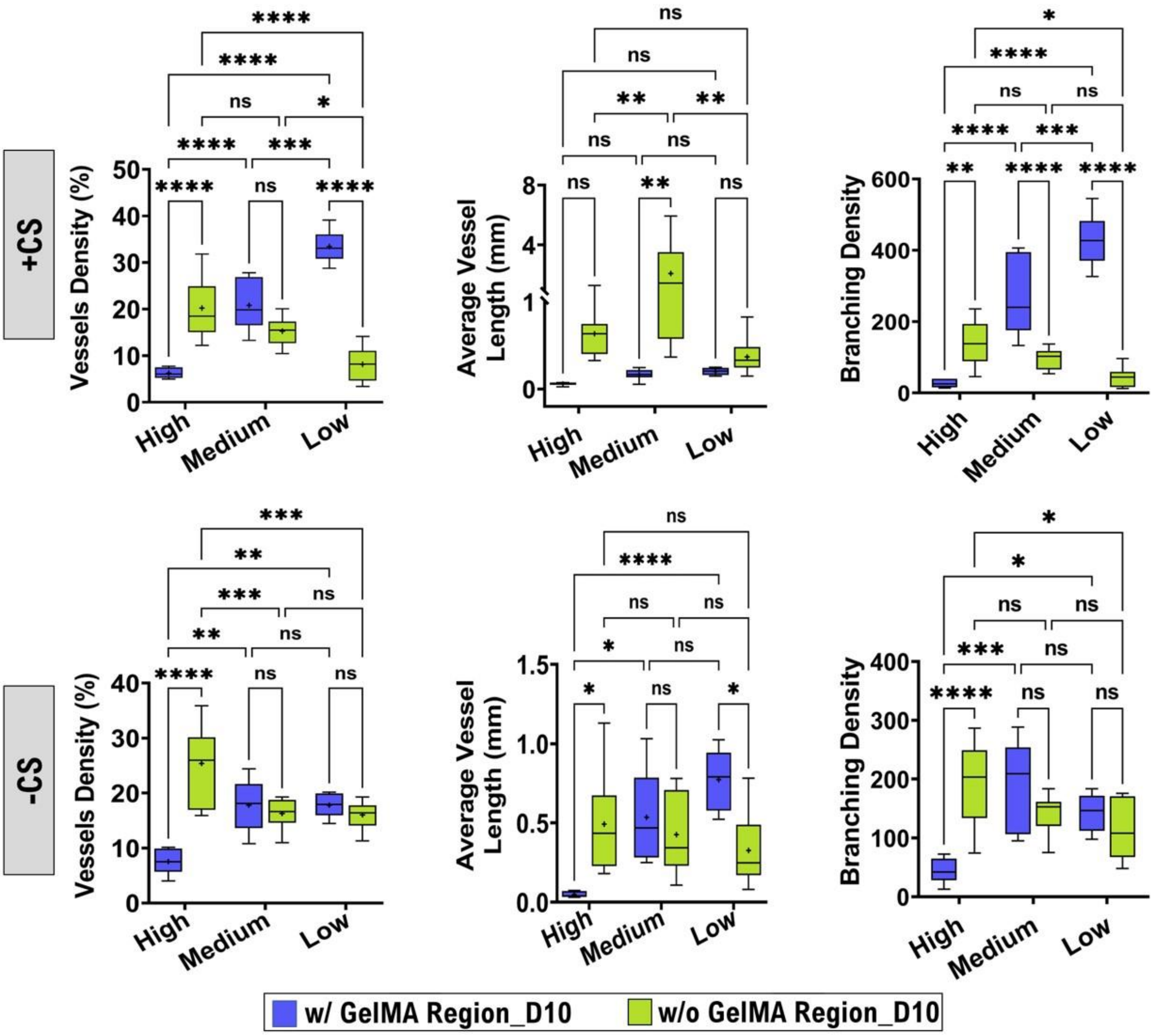


***Figure S7. Influence of bioprinted neighboring GelMA regions on controlling vascular network properties within bioprinted aptamer regions in the presence (or absence) of CS (added on D4) on D10.*** *For comparison, the parallel line designs with high, medium and low spatial resolutions printed with neighboring GelMA regions are referred as "w/ GelMA Region_D10" samples and the vascular tree design samples bioprinted only with programmable bioink are referred as "w/o GelMA Region_D10" samples. Among these samples, the vascular properties within the printed aptamer regions were compared based on the printed line widths. Comparative analysis of the representative vascular network properties such as (A) vessel density (%) and (B) average vessel length (mm) in the presence/absence of CS (added on D4) at different printed line widths on D10.*

### Supplementary Tables & Videos

**Table S1.** Aptamer sequences and their characteristics were used in the present study. Tm denotes the melting temperature (50 mM NaCl), MW is molecular weight and N signifies the number of nucleotides.

| Aptamer | Sequence (5'→ 3') | Tm | MW | N |
|---|---|---|---|---|
| **VEGF Specific Aptamer** | /5Acryd/CGA TCG TAT CAG TCC ACA AGC CCG TCT TCC AGA CAA GAG TGC AGG GC | 70.8 °C | 14665.6 | 47 |
| **Complementary Sequence (CS)** | CGC CCT GCA CTC TTG TCT GGA AGA CGG GCT TGT GGA CTG ATA CGA TCG | 71.3 °C | 14791.6 | 48 |
| **Fluorescently labelled CS ($CS_F$)** | /5Alexa488N/CGC CCT GCA CTC TTG TCT GGA AGA CGG GCT TGT GGA CTG ATA CGA TCG | 71.3 °C | 15487.2 | 48 |

**Table S2.** Parameters used for the reaction-diffusion model of free VEGF release within aptamer functionalized hydrogels.

| Parameters | Values |
|---|---|
| VEGF concentration in Aptamer region at $t$ = 0hr $[\boldsymbol{C_{VEGF-aptamer}}]$ | 0.25 nM |
| VEGF concentration in GelMA region at $t$ = 0hr $[\boldsymbol{C_{VEGF-GelMA}}]$ | 0.012 nM |
| VEGF Diffusivity in the absence of CS $[\boldsymbol{D_{VEGF-CS}}]$ | 0 $m^2/s$ |
| VEGF Diffusivity in the presence of CS $[\boldsymbol{D_{VEGF+CS}}]$ | 1.8 x $10^{-13}$ $m^2/s$[5] |

**Video S1.** 3D Bioprinting of high-resolution line design using both GelMA and programmable bioinks simultaneously.

**Video S2.** In-silico analysis of free VEGF diffusion from aptamer region to GelMA region in the absence of CS during 240 hr of experiment, for high-resolution design.

**Video S3.** In-silico analysis of free VEGF diffusion from aptamer region to GelMA region in the presence of CS during 240 hr of experiment, for high-resolution design.

**Video S4.** In-silico analysis of free VEGF diffusion from aptamer region to GelMA region in the absence of CS during 240 hr of experiment, for medium-resolution design.

**Video S5.** In-silico analysis of free VEGF diffusion from aptamer region to GelMA region in the presence of CS during 240 hr of experiment, for medium-resolution design.

**Video S6.** In-silico analysis of free VEGF diffusion from aptamer region to GelMA region in the absence of CS during 240 hr of experiment, for low-resolution design.

**Video S7.** In-silico analysis of free VEGF diffusion from aptamer region to GelMA region in the presence of CS during 240 hr of experiment, for low-resolution design.

#### References


[1] D. Rana, A. Kandar, N. Salehi-Nik, I. Inci, B. Koopman, J. Rouwkema, *Bioact Mater* **2022**, *12*, 71.

[2] D. Rana, P. Padmanaban, M. Becker, F. Stein, J. Leijten, B. Koopman, J. Rouwkema, *Mater Today Bio* **2023**, *19*, 100551.

[3] J. W. Nichol, S. T. Koshy, H. Bae, C. M. Hwang, S. Yamanlar, A. Khademhosseini, *Biomaterials* **2010**, *31*, 5536.

[4] S. Bertlein, G. Brown, K. S. Lim, T. Jungst, T. Boeck, T. Blunk, J. Tessmar, G. J. Hooper, T. B. F Woodfield, J. Groll, et al., *Advanced Materials* **2017**, *29*, 1703404.

[5] L. Abune, N. Zhao, J. Lai, B. Peterson, S. Szczesny, Y. Wang, *ACS Biomater Sci Eng* **2019**, DOI 10.1021/ACSBIOMATERIALS.9B00423/SUPPL_FILE/AB9B00423_SI_001.PDF.
