## Supplementary figures and images for "Bioprinting of aptamer-based programmable bioinks to modulate multiscale microvascular morphogenesis in 4D"

### Video S2

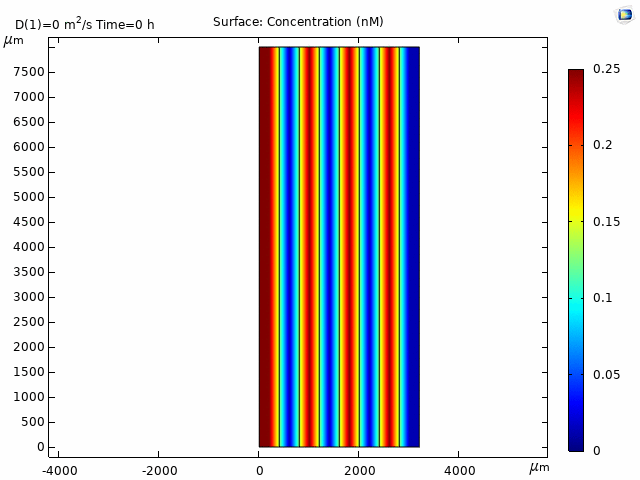

### Video S3

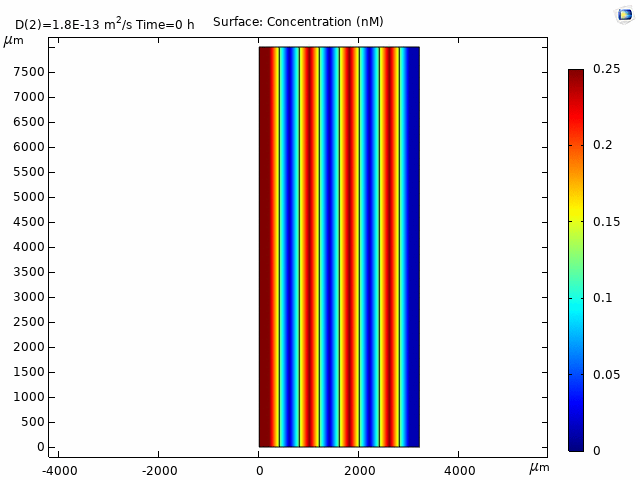

### Video S4

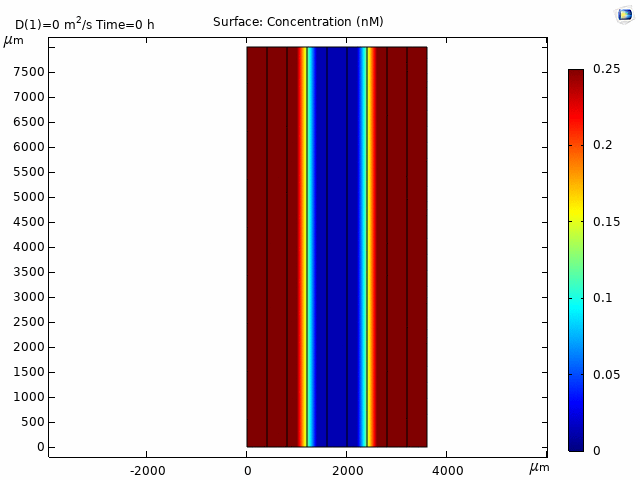

### Video S5

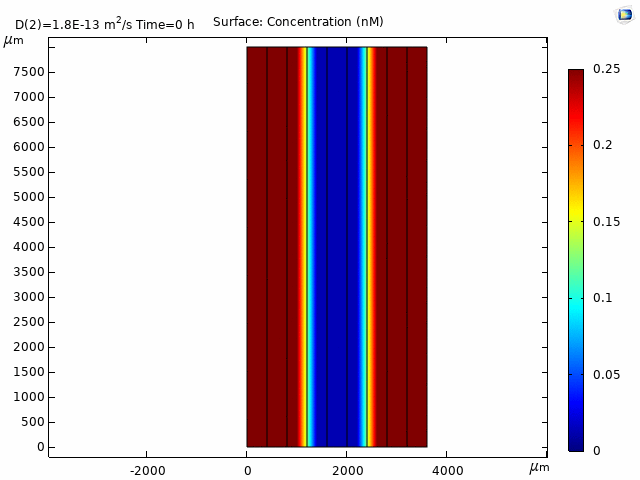

### Video S6

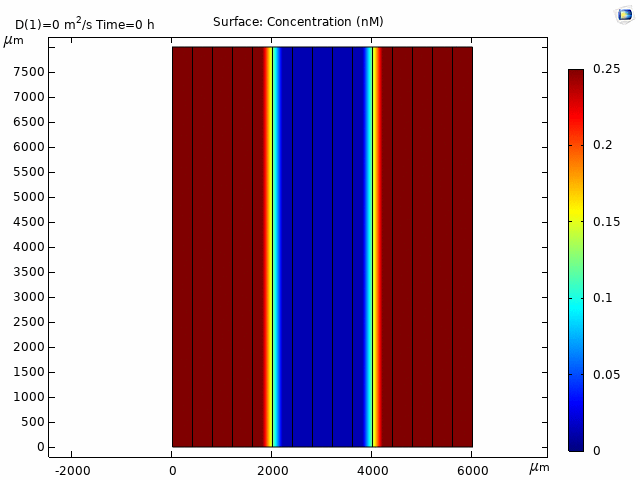

### Video S7

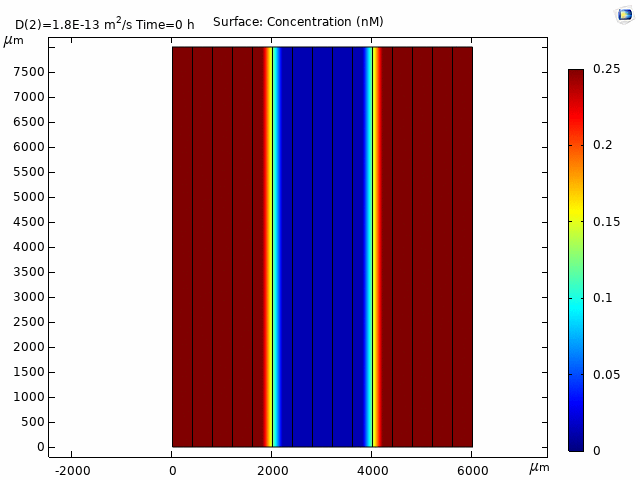
